# Realistic Coupling enables flexible macroscopic traveling waves in the mouse cortex

**DOI:** 10.1101/2025.07.05.663278

**Authors:** Guanhua Sun, James Hazelden, Ruby Kim, Daniel B. Forger

**Author notes:** These authors contributed equally to this work (Guanhua Sun, James Hazelden). Department of Applied Mathematics, University of Washington, United States (James Hazelden).

## Abstract

Traveling waves are ubiquitous in neuronal systems across different spatial scales. While microscopic and mesoscopic waves are relatively well studied, the emergence of macroscopic traveling waves remains less understood. Here, by modeling the mouse cortex using spatial transcriptomic and connectivity data, we show that realistic cortical connectivity can generate a significantly higher level of macroscopic traveling waves than artificial local and uniform connectivity across multiple oscillation frequency bands, with the strongest advantage appearing in the theta, alpha, and beta frequency bands. By probing the model in different dynamic regimes, we find that macroscopic wave activity depends on both network connectivity and excitatory coupling strength, with a non-monotonic dependence on coupling. Together, our work shows how flexible macroscopic traveling waves can emerge in the mouse cortex and offers a computational framework to further study traveling waves in the mouse brain at the single-cell level.

## Introduction

Traveling waves have emerged as a central topic among different large-scale spatiotemporal dynamics observed in neuronal systems Muller et al. (2018). These waves are observed at different frequencies during various brain activities, ranging from slow-wave activity Massimini (2004); Liang et al. (2021), sleep spindles Muller et al. (2016), to faster oscillations in alpha Zhang et al. (2018); van Kerkoerle et al. (2014), beta Bhattacharya et al. (2022); Rubino et al. (2006), and gamma van Kerkoerle et al. (2014); Aggarwal et al. (2022) frequency bands, as well as in broadband (5-40 Hz) activity Davis et al. (2020). These waves also appear in various brain regions across species, including the hippocampus Lubenov and Siapas (2009); Patel et al. (2012); Zhang and Jacobs (2015), visual cortex Sato et al. (2012), prefrontal cortex Bhattacharya et al. (2022), motor cortex Rubino et al. (2006), or span across multiple brain regions Liang et al. (2023). More importantly, these waves have long been conjectured to regulate brain functions through mechanisms such as synaptic plasticity Ermentrout and Kleinfeld (2001); Muller et al. (2018). And researchers have recently found that traveling waves can modulate important brain functions such as memory, visual processing, and sleep Mohan et al. (2024); Aggarwal et al. (2022); Miyamoto et al. (2017).

The prevalence and important implications of traveling waves have prompted researchers to investigate their causes and organization. In these studies, computational modeling has been useful due to its flexibility. Researchers have developed both microscopic models Davis et al. (2020); Naze et al. (2015); Ermentrout and Kleinfeld (2001) and mesoscopic models Roberts et al. (2019); Liang et al. (2023) to explore the mechanical properties and computational implications of traveling waves. However, most of these studies focus on microscopic or mesoscopic waves that appear in a single brain region. Large-scale, macroscopic waves across multiple brain regions have been less studied. Moreover, these models are usually constructed with local or uniform connectivity and lack the realistic geometry and connectivity of the brain. It has been shown that in a coupled neuronal network, the coexistence of global wave activity with locally asynchronous states only arises when the number of oscillators is high enough, where local and global activities can coexist Davis et al. (2021). At the same time, it has been argued that modeling neuronal networks at single-cell resolution will yield more precise discoveries than the mean-field model Dura-Bernal et al. (2024); Deco et al. (2008). Therefore, it is ideal to study traveling waves in a large domain such as the cortex at single-cell resolution.

However, there are two significant challenges: First, a microscopic wiring diagram of the mouse brain at the whole-brain level is not available. But we do have mesoscopic connectivity data provided by the Allen Brain Atlas Oh et al. (2014); Knox et al. (2018); Harris et al. (2019), which will be used in this work. Second, simulating and measuring wave activity in a large 3D computational domain with realistic brain geometry is computationally expensive. Simulations of large-scale neuronal networks with realistic neuronal models are usually conducted on supercomputers Yamaura et al. (2020); Markram et al. (2015).

In this work, we overcome these two challenges in order to study the emergence and propagation of macroscopic traveling waves in the mouse cortex. First, we propose a network-generating algorithm that integrates a spatial transcriptomic dataset that provides the detailed spatial and molecular profile of neurons Zhang et al. (2023), with voxelized connectivity data Knox et al. (2018) of the mouse brain. We use this algorithm to create a neuron-to-neuron connectome for one hemisphere of the mouse cortex, which we call *Allen* connectivity. We then constructed a 3D computational model with both excitatory and inhibitory neurons on top of the Allen connectivity. To overcome the second difficulty, we built our own parallel computational framework that can simulate the constructed model with a desktop GPU.

We simulate this model under stochastic layer-4 input, and we quantify macroscopic wave activity with a three-dimensional phase-gradient analysis. Across a broad parameter sweep, Allen connectivity produces stronger macroscopic waves than local or uniform connectivity, especially in the theta, alpha, and beta bands. We further find that wave activity depends non-monotonically on excitatory coupling strength and is closely linked to network synchrony. Together, these results show how realistic cortical connectivity can support flexible macroscopic traveling waves and provide a computational framework for studying cortex-wide wave dynamics at single-cell resolution.

## Results

### Construction of the cortical connectivity

A recent spatial transcriptomic dataset (Zhuang-ABCA1) Zhang et al. (2023) provides detailed positions of neurons in the mouse brain along with their molecular properties, including the neurotransmitter information (Fig. 1a, Use of the spatial transcriptomic data). To build a computational model using this data, we need to connect neurons in Zhuang-ABCA1 based on the realistic connectivity of the mouse brain that has been experimentally measured. Previously, Oh et al. (2014) and Harris et al. (2019) have provided a mesoscopic connectivity matrix, known as the “projectome”, which is derived from thousands of viral tracing experiments to measure the strength of connections between different brain regions mesoscopically. This projectome was later refined to a finer level using a statistical model introduced by Knox et al. (2018), where the strength of connections is provided on a voxel (100*μm*^3^) resolution. Because both neuronal (Zhuang-ABCA1) and the voxelized connectivity data are registered with the Allen Brain Atlas Common Coordinate Framework (ABA-CCF) Wang et al. (2020), we now introduce an algorithm that integrates the two datasets to construct a neuron-to-neuron connectivity.

**Figure 1.**
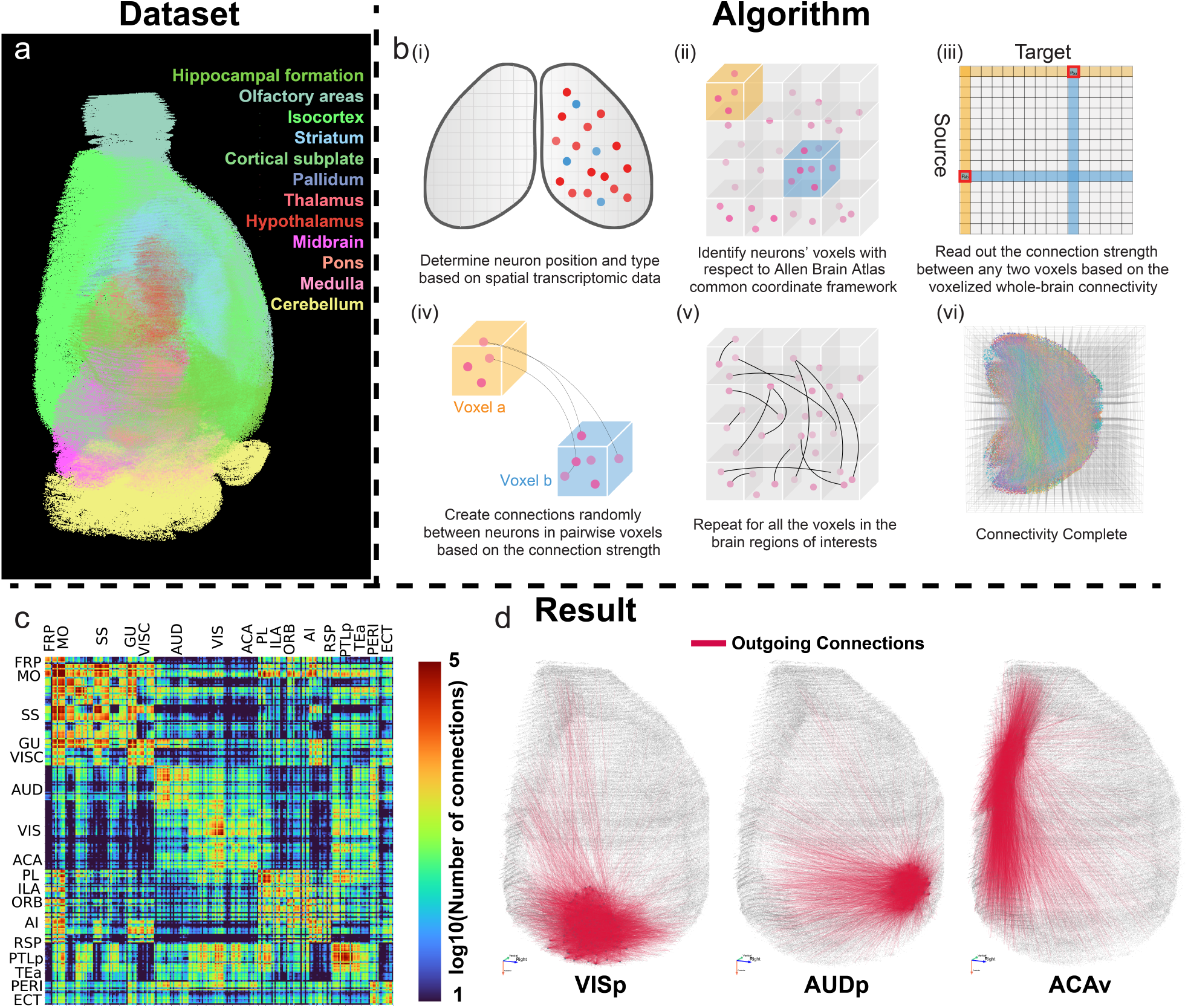
Construction of the cortical connectivity. **(a)** Visualization of all neurons in the hemispherical dataset we use to build the cortical connectivity. The hemisphere contains about 1 million neurons in total, including about 300,000 cortical neurons. **(b)** Algorithm of building the connectome: i) First, we identify neuronal excitatory/inhibitory information based on the spatial transcriptomic data. ii) We then identify the voxelized position for all the neurons. iii) Based on the voxelized projection data, we calculate the number of connections between every pair of voxels (see Use of the voxelized connectivity data). iv) Connections are created randomly between the neurons in each pair of voxels. v) Repeat for all voxels in regions of interest. vi) Complete the connectivity. **(c)** The connectivity matrix shows the number of connections between different cortical regions(in log 10 scale). A full version of the connectivity matrix with annotated regions is shown in figure Supplement 1. **(d)** Visualization of outgoing connections from sampled neurons in the primary visual cortex (VISp, left), primary auditory cortex (AUDp, middle), and the ventral anterior cingulate areas (ACAv, right).

Consider constructing a connectivity for N neurons that are contained in M voxels, and let *X*_*i*_ represent the physical position of the ith neuron. We have access to the voxelized projection matrix W, where *W*_*ab*_ denotes the projection strength from voxel a to voxel b. Our connectivity algorithm requires only a single density parameter, ρ, which determines the average number of connections per neuron, thereby controlling the network’s sparsity (Use of the voxelized connectivity data). Using this information, we propose the following algorithm to construct the connectivity for the target neurons (Fig. 1b and Methods):

**Algorithm 1.**
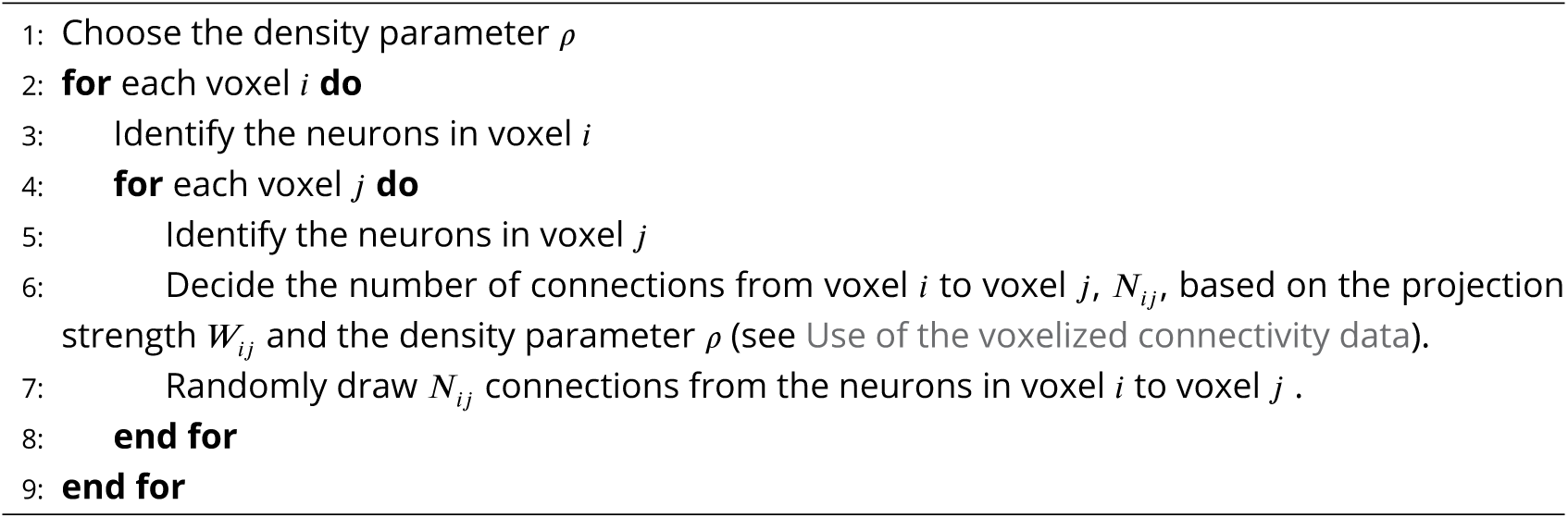
Construction of cortical connectivity.

Applying this algorithm to all cortical neurons identified in Zhuang-ABCA1, we obtain a cortical connectivity that consists of 297,812 neurons, and we choose each neuron to have 300 connections on average, resulting in a network sparsity of around 0.1 % (Construction and storage of the connectivity). The cortical connectivity we constructed is highly heterogeneous, where the number of connections between different regions can vary across four orders of magnitude (Fig. 1c). At the same time, both the in-degree and out-degree distribution of the connectome follow a realistic log-normal distribution Buzsáki and Mizuseki (2014) (figure Supplement 1). Since the complete connectome is difficult to visualize, we show some connections from different cortical regions, including the primary visual, auditory cortex, and the ventral anterior cingulate area (Fig. 1d).

### Macroscopic traveling waves emerge from random stimulation through realistic connectivity

Equipped with the Allen connectivity, we built a single-cell-resolution cortical model to test whether realistic anatomical wiring is sufficient to convert a generic stochastic input into a coordinated, macroscopic traveling wave. Each neuron is a single-compartment Hodgkin-Huxley unit Pospischil et al. (2008), with its excitatory (glutamatergic, AMPA) or inhibitory (GABAergic) identity inherited from the Zhuang-ABCA-1 transcriptomic labels, and synapses are conductance-based with first-order kinetics (Neuronal and synaptic model). The full hemispheric network of 297,812 neurons is simulated on a single GPU (Simulation).

Spiking networks of this scale can also generate self-sustained activity in similar regimes Vogels and Abbott (2005); Kumar et al. (2008), which differs fundamentally from externally imposed noise Destexhe and Contreras (2006), and traveling waves have been reported under such conditions Davis et al. (2021). Therefore, to mimic the stochastic drive that cortex receives from subcortical regions like thalamus, we deliver a 10 Hz Poisson spike train to layer-4 excitatory neurons (Figure 2a), since layer 4 is the canonical thalamocortical input layer Rockland (2019); each Poisson event applies a fixed voltage bump *V*_stim_, and no other external input is applied.

**Figure 2.**
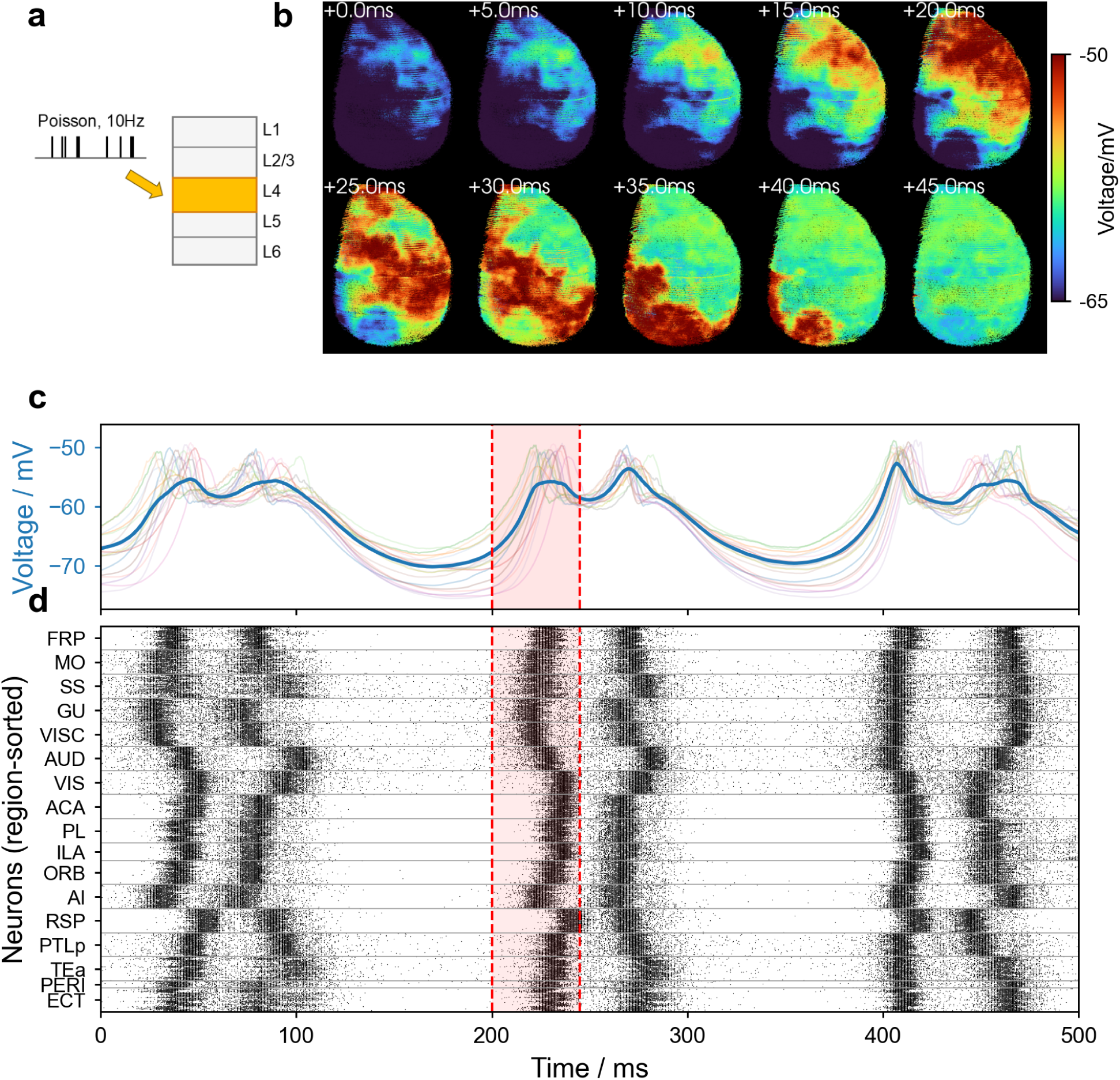
Macroscopic traveling waves emerge in response to random layer-4 stimulation through Allen connectivity. **(a)** Stimulation protocol: a 10 Hz random Poisson spike train delivers a fixed-amplitude voltage bump *V*_stim_ to all layer-4 excitatory neurons; no other external input is applied. **(b)** Two rows of five spatial snapshots of the local-mean (KNN = 300) intracellular voltage, sampled every 5ms across the window highlighted in **(c)**. A coherent wavefront propagates along the anterior-to-posterior axis. **(c)** Global mean intracellular voltage of all neurons (dark blue) overlaid on per-region mean voltages (light traces). The red dashed lines mark the snapshot window visualized in **(b)**. **(d)** Region-sorted raster plot of ∼40,000 neurons sampled evenly from major cortical regions, anterior-to-posterior. See Videos for the movie of the simulation.

Under this protocol, we immediately observe macroscopic traveling waves emerge across the cortex (**Figure 2** and Videos). The global mean voltage and the region-sorted raster (Figure 2c, d) reveal oscillatory activity that is well synchronized across regions, while the local-mean intracellular voltage maps over a representative 50 ms window (Figure 2b) reveal a coherent wavefront sweeping along the anterior-posterior axis, consistent with previously reported cortex-wide waves Massi-mini (2004); Liang et al. (2021); Aggarwal et al. (2024). The corresponding single-neuron-resolution view of the same simulation, with no spatial averaging, is shown in figure Supplement 1.

### Realistic connectivity produces stronger macroscopic waves than local or uniform connectivity

In past modeling studies of traveling waves, it is common to adopt uniform-random, local, or a combination of the two kinds of connectivity. Therefore, we now simulate three models differing in connectivity in order to compare and investigate whether such macroscopic waves are presented in **Figure 2** only emerge from the Allen connectivity. For comparison, we constructed two additional types of connectivity that are common in past studies of traveling waves:

1. Uniform connectivity: Neurons are connected across the entire cortex randomly (Fig. 3b).
2. Local connectivity: Neurons are connected to a fixed number of their closest neighbors (Fig. 3c).

**Figure 3.**
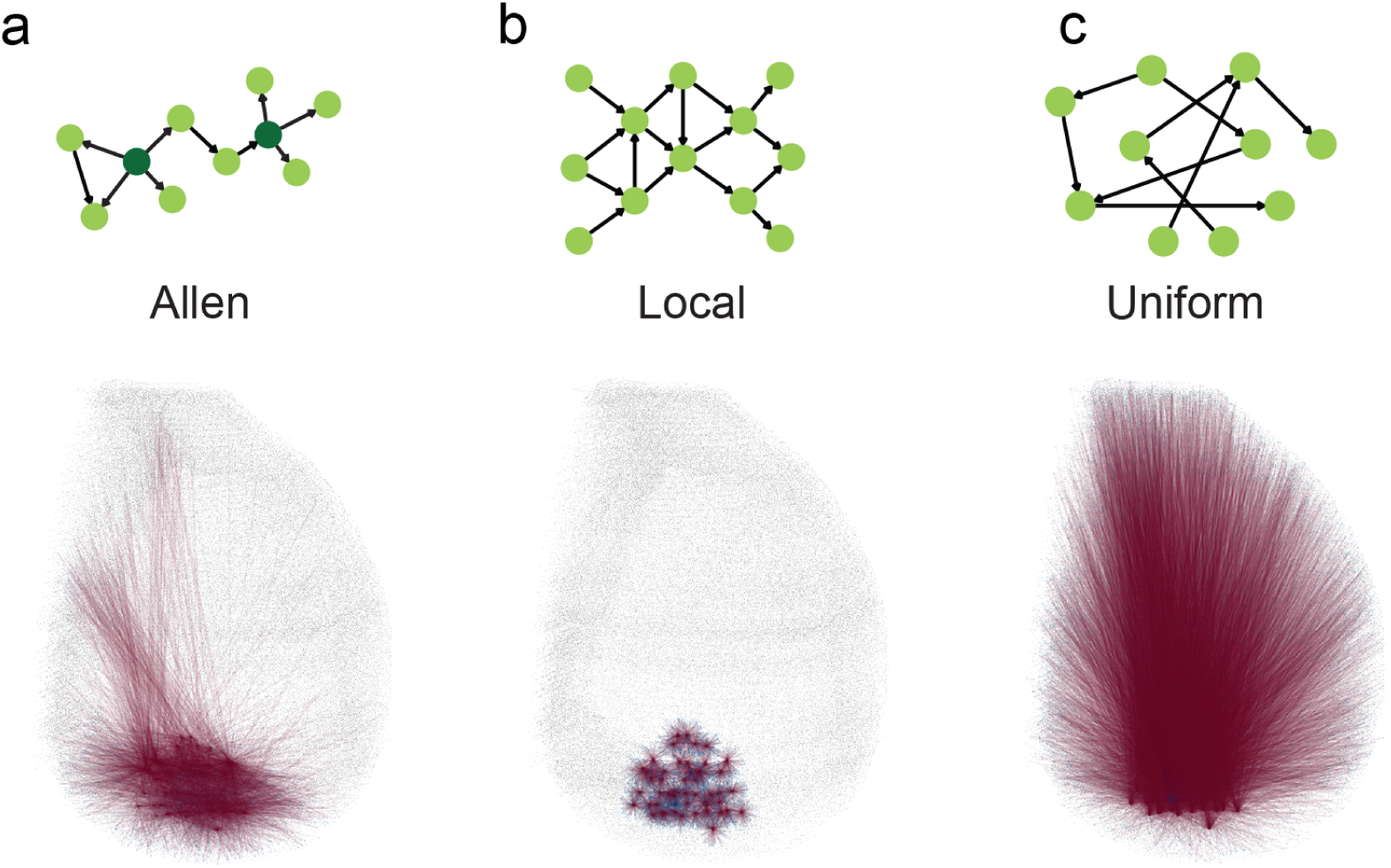
Top: schematics of Allen **(a)**, local **(b)**, and uniform connectivity **(c)**; bottom: Visualization of outgoing connections from the same 50 randomly selected neurons in the primary visual cortex (VISp) for the three connectivity.

Here we visualize the outgoing connections from 50 randomly selected neurons in the primary visual cortex to show the difference between the three connectivity. Under Allen connectivity, the outgoing connections project to different regions heterogeneously (Fig. 3a); with local connectivity, the outgoing connections only reach around the adjacent areas where the sampled neurons are located (Fig. 3b); and with uniform connectivity, the outgoing connections tend to cover the whole cortex evenly (Fig. 3c). Although the topology of three connectivity is different, they share the same out-degree distribution based on the way they are constructed (see Construction and storage of the connectivity).

To properly quantify the traveling waves observed in simulations, we applied a phase-based method from image processing Fleet and Jepson (1990), which has been adopted to analyze neuronal dynamics Rubino et al. (2006), and extended it to 3-D data. We first interpolate the simulated voltage onto a regular Cartesian grid that follows the Allen Brain Institute common coordinate framework (CCFv3); the voltage signal is then band-pass filtered into a narrow frequency band, and the Hilbert transform is applied to the narrowband-filtered grid signal to extract the band-specific generalized phase. We then compute the gradient field of the phase signal on the grid. The phase gradient directionality (PGD), an indicator of planar wave activity Rubino et al. (2006), is then used as our quantitative measure of macroscopic wave activity (see Quantitative measurement of neuronal activity for the full pipeline).

In **Figure 4** and Videos, we compare the PGD produced by three representative simulations with different connectivity but the same parameters otherwise. The phase snapshots in **Figure 4** span a 20 ms window centered on the PGD peak. Although both Allen and local connectivity produce visible wave activity, Allen connectivity produces waves that are much more macroscopic, while local connectivity produces locally organized planar waves, the form most frequently reported in previous experimental studies Lubenov and Siapas (2009); Bhattacharya et al. (2022) and computational models with local connectivity Keane and Gong (2015). Uniform connectivity does not generate any apparent structure. This qualitative observation is reflected in the PGD time series (Figure 4b; analogous comparisons in the delta, alpha, beta and gamma bands are shown in figure Supplement 2, figure Supplement 3, figure Supplement 4 and figure Supplement 5 respectively): Allen connectivity reaches the highest PGD in this representative simulation, with local and uniform remaining markedly lower.

**Figure 4.**
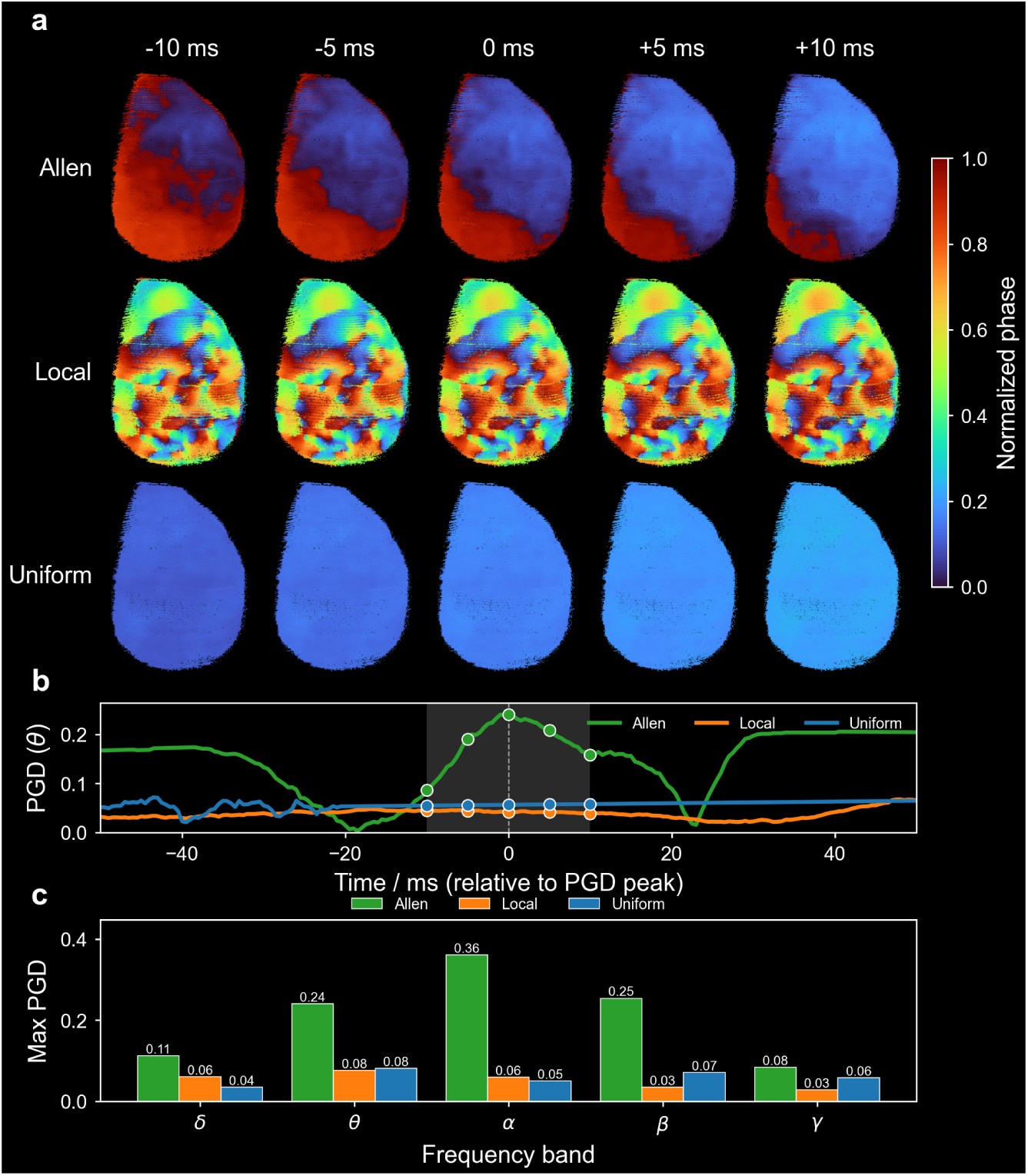
Allen connectivity produces a higher level of macroscopic wave activity than local and uniform connectivity. **(a)** Theta-band (4-8 Hz) phase snapshots from three representative simulations using Allen (top), local (middle) and uniform (bottom) connectivity under matched Poisson stimulation, sampled at −10, −5, 0, +5 and +10ms relative to the peak of phase gradient directionality (PGD). Colors represent the theta-band generalized phase of each neuron, normalized between 0 and 1. **(b)** Theta-band PGD of the three simulations versus time, aligned to the PGD peak. **(c)** Per-band comparison of the maximum PGD of the three representative simulations shown in **(a)**, one bar per band per connectivity. Please see Videos for animations and Quantitative measurement of neuronal activity (illustrated in figure Supplement 1) for the calculation of PGD.

### Network coupling and synchrony jointly shape macroscopic waves across the parameter plane

Traveling waves are known to propagate differently under different dynamical states, even when the underlying connectivity is fixed Muller et al. (2018). To assess whether this connectivity-dependent advantage holds beyond a single set of parameters, we systematically swept the coupling strength *g*_AMPA_ and the Poisson stimulus magnitude *V*_stim_ on a 10 × 12 grid for each connectivity (Figure 5a, Simulation). Averaged over the full grid, Allen connectivity continues to produce the highest per-band PGD relative to local and uniform connectivity (Figure 5b), showing that the advantage seen Figure 4c is not specific to one set of parameters. We do note that for the delta band, local and Allen connectivity are able to generate a similar level of strong macroscopic wave activities. The same parameter plane also reveals three qualitatively distinct dynamic regimes for Allen connectivity (Figure 5a): coherent macroscopic waves at weak coupling, a disordered (asynchronous irregular) state at medium coupling in which wave structure collapses, and a globally synchronous regime at strong coupling that again supports waves.

**Figure 5.**
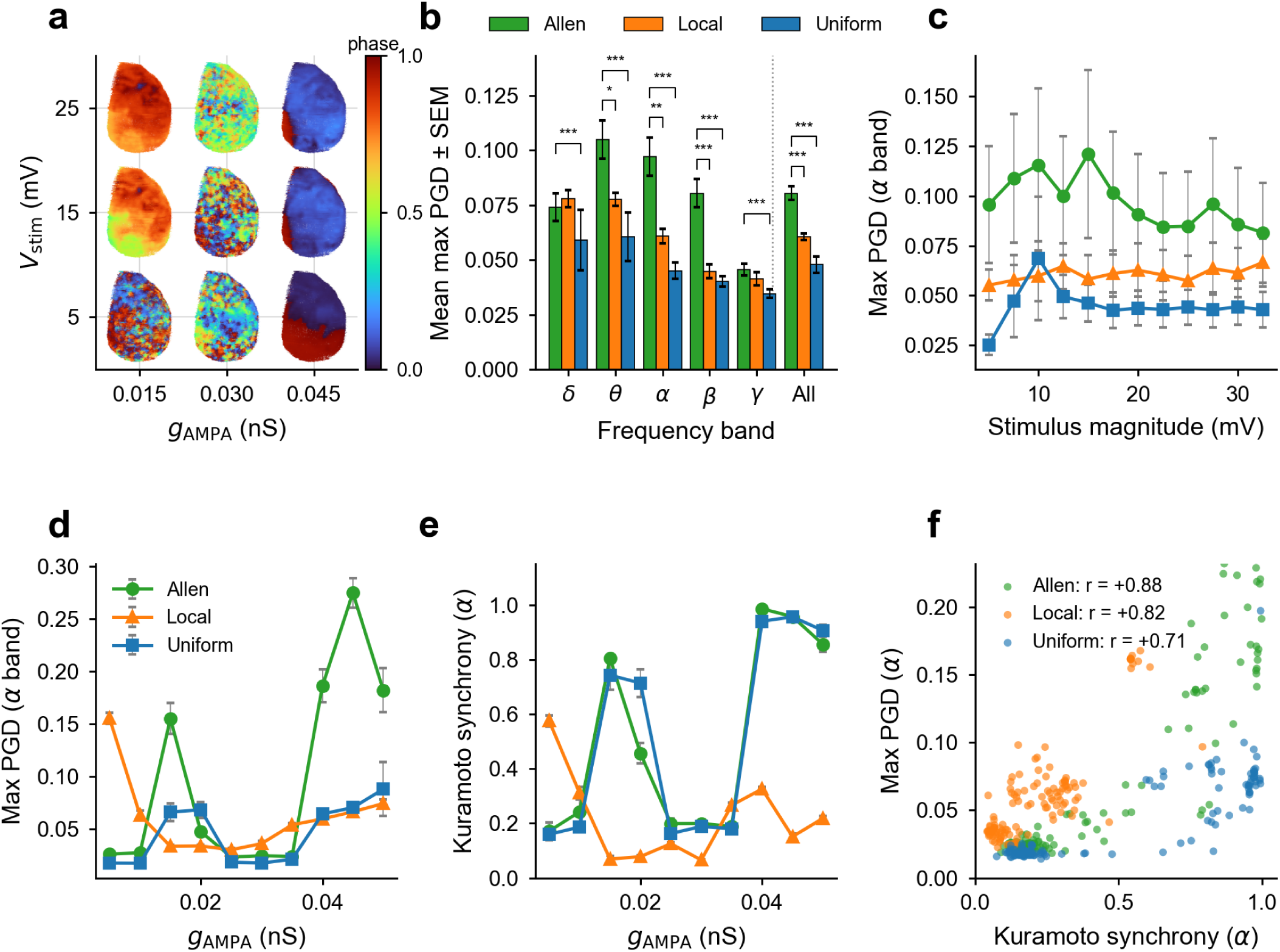
Coupling strength, connectivity, and network synchrony jointly shape macroscopic-wave activity. **(a)** Schematic of the three dynamic regimes that emerge across the (*g*_AMPA_, *V*_stim_) parameter plane for Allen connectivity. At weak-to-intermediate coupling, the network supports *coherent* macroscopic traveling waves; at medium coupling, increased synaptic drive pushes the network into a *disordered* (asynchronous irregular) state where wave structure collapses; at strong coupling, the network reorganizes into a *globally synchronous* regime that again supports wave propagation. Insets show representative phase snapshots from each regime. **(b)** Per-band max PGD (mean ± SEM across the Poisson *g*_AMPA_ × *V*_stim_ grid), plus the across-bands aggregate (‘All’), for Allen, local and uniform connectivity, with significance stars; full per-band (*g*_AMPA_, *V*_stim_) heatmaps in figure Supplement 6. **(c)** Max PGD (alpha band) versus stimulus magnitude, error bars across *g*_AMPA_. **(d)** Max PGD (alpha band) versus excitatory conductance *g*_AMPA_, error bars across *V*_stim_.**(e)** Kuramoto order parameter versus *g*_AMPA_ in the alpha band. **(f)** Synchrony-PGD scatter, one point per (*g*_AMPA_, *V*_stim_) combination, with Pearson *r* per connectivity (Allen *r*= 0.88, local *r*= 0.82, uniform *r*= 0.71 in alpha; all bands positive and significant at *p <* 10−3). Per-band versions of (c)-(e) are shown in figure Supplement 1, and per-band synchrony-PGD scatters in figure Supplement 2.

To dissect the non-monotonic coupling response that distinguishes the three regimes, we examine how PGD varies along each axis of the (*g*_AMPA_, *V*_stim_) plane in the alpha band - the band that carries a large advantage for Allen connectivity (Figure 5b). Macroscopic wave activity is relatively insensitive to the stimulus magnitude (Figure 5c) but depends strongly and non-monotonically on the coupling strength (Figure 5d): for Allen connectivity, PGD peaks at weak coupling, collapses through a medium-coupling window (*g*_AMPA_ ≈ 0.020-0.035 nS), and recovers at strong coupling. The uniform connectivity also follows this non-monotonic relationship, but has much lower max PGD at peaks compared to Allen connectivity. Local connectivity follows a different trajectory, peaking at weak coupling, then decaying almost monotonically with *g*_AMPA_. These trends hold band by band (figure Supplement 1): the peak-dip-recovery profile is visible in every band for Allen and uniform, while local decays almost monotonically across all bands.

To diagnose the mechanism behind this profile, we measured the Kuramoto order parameter *R*(*t*) = *e*^*i*Φ(*x,t*)^ _*x*_ from the generalized phase field Φ(*x*, *t*) of each frequency band and recorded its maximum over each simulation window (Quantitative measurement of neuronal activity). The resulting synchrony curve (Figure 5e) resembles the trend of the maximum PGD well (Figure 5d). For Allen connectivity in the alpha band, mean synchrony peaks at weak coupling, collapses in the medium-coupling window, and recovers at strong coupling. Across the full 120-point (*g*_AMPA_, *V*_stim_) grid, synchrony and PGD are positively correlated in every band and every connectivity (Pearson *r* = 0.30-0.92, all *p* < 10^−3^; Figure 5f, with per-band scatters in figure Supplement 2). Therefore, the PGD trough at medium coupling may be a synchrony trough: increased synaptic drive pushes the network into an asynchronous irregular state, while strong coupling reestablishes rhythmic synchronization that supports wave propagation. This synchrony-to-wave bottleneck seems more significant to the networks with long-range connectivity (Allen, uniform), partly because long-range connections can augment synchrony in the coupled neuronal network Bazhenov et al. (2008). We also note that although uniform connectivity is able to achieve almost an identical level of synchrony to that of Allen connectivity, the PGD remains much lower due to the loss of spatial organization within.

### The macroscopic wave findings extend to the local field potential estimate

Across the paper, our analyses use the intracellular voltage as the primary measurement for the PGD. To verify that the macroscopic wave structure we identify is also visible in a quantity comparable to experimentally recorded local field potentials (LFP), we followed a post-hoc pipeline Telenczuk et al. (2020) that reconstructs the net synaptic current received by every neuron directly from the voltage traces and the simulation’s connectivity (LFP estimation from intracellular voltage). The same 3-D phase-gradient pipeline is then applied to the reconstructed LFP. The propagating wavefront seen in the local-average voltage is recovered, with cleaner contrast, in the LFP estimate (Figure 6a; Videos). Voltage and LFP max PGD are tightly correlated across the parameter grid (Pearson *r* = 0.78-0.89 per band, *p* < 10^−25^ for every band; Figure 6b), and the LFP version carries roughly 2-3 × more PGD than the underlying voltage signal (paired *p* < 10^−13^ in every band; Figure 6c). Most importantly, Allen connectivity remains the highest-PGD connectivity in every band when measured on the LFP signal (Figure 6d), and the non-monotonic coupling profile from **Figure 5** is preserved (Figure 6e). These results suggest that the macroscopic-wave findings are not specific to intracellular voltage and can also be recovered in an LFP-like signal.

**Figure 6.**
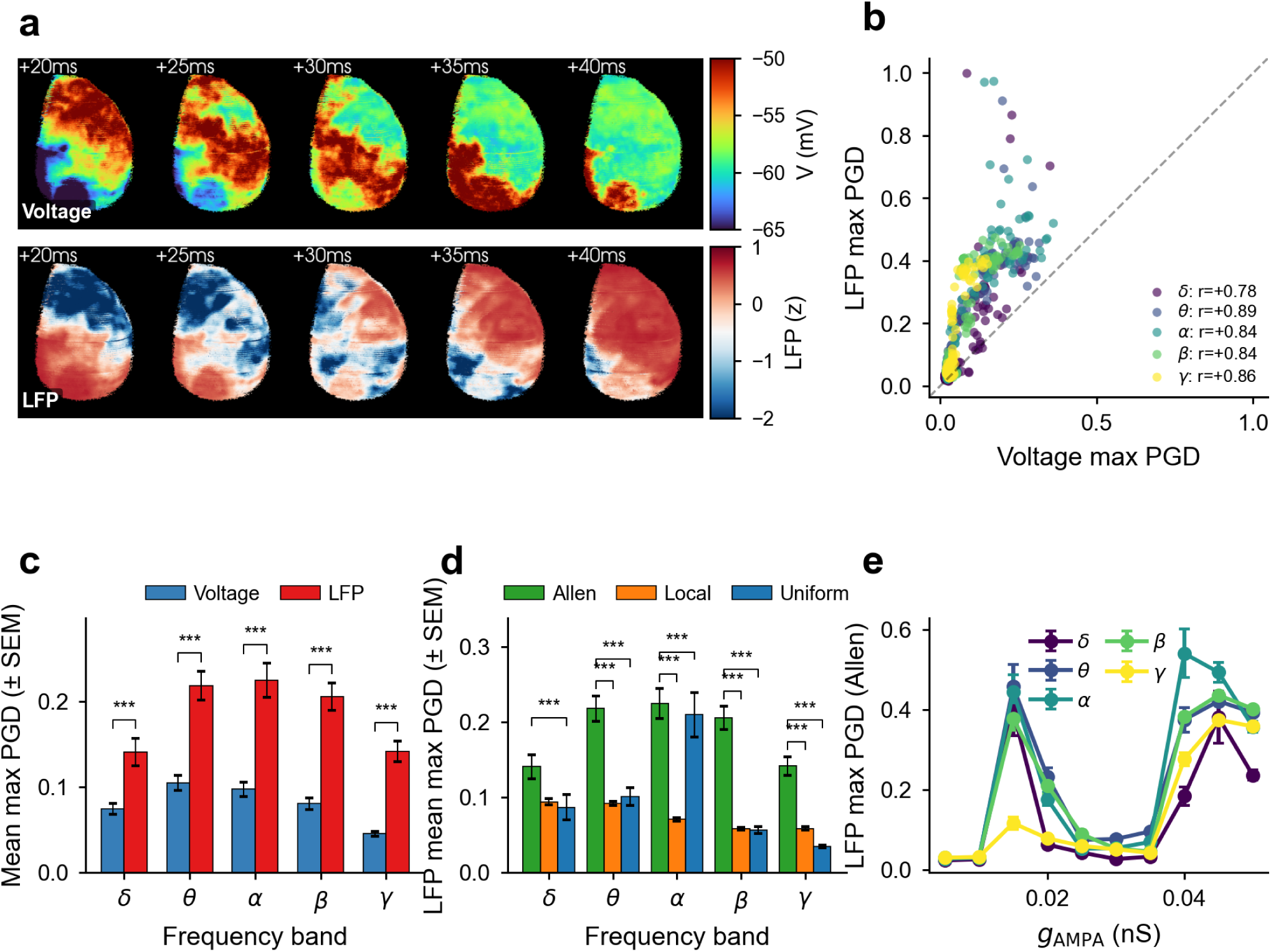
Macroscopic waves are maintained by the local field potential estimate. (a) Paired snapshots of the local-average voltage (top, clipped to [−65, −50] mV) and the synaptic-LFP estimate (bottom, *z*-scored) across five consecutive 5-ms frames in a representative Allen Poisson simulation. The same propagating wavefront sweeps across the cortex in both modalities. (b) Voltage versus LFP max PGD scatter for Allen connectivity, one point per (*g*_AMPA_, *V*_stim_, band); Pearson *r* = 0.78-0.89 per band, all *p* < 10^−25^. (c) Mean max PGD per band, voltage versus LFP, for Allen connectivity (paired *p* < 10^−13^ for every band; LFP carries ≈ 2-3× higher PGD than the intracellular voltage). (d) LFP-based mean max PGD per band, three connectivities; Allen wins in every band against both local and uniform. (e) LFP max PGD versus *g*_AMPA_ for Allen connectivity, one line per band; the same three-regime profile (peak, dip, recovery) seen in voltage is preserved in LFP. See LFP estimation from intracellular voltage for the LFP estimation pipeline.

## Discussion

### Construction of the Allen connectivity

In this work, we present an algorithm that integrates the spatial transcriptomic dataset, which provides the spatial distribution and molecular properties of neurons across the mouse brain, and the mesoscopic connectivity dataset that offers the connectivity information between brain regions on a voxelized level. The algorithm is efficient and can be easily adapted for similar datasets of the mouse brain or for even other species. In essence, our algorithm belongs to the family of stochastic block models Holland et al. (1983), where the block structure is given by the voxelization of the Allen Brain Atlas and the inter-block connection probabilities are set by the voxelized projection strengths. The projectome Knox et al. (2018) we use to build the Allen connectivity is derived from a series of viral-tracing experiments that reflect the connection strength between different regions Oh et al. (2014). However, the relationship between the actual connectome and the mesoscopic projectome remains unclear, and our current model inherits several important simplifications from this dataset.

First, the voxelized projection data is not cell-type specific. In our model, we distinguish only two neuronal populations: glutamatergic (excitatory) and GABAergic (inhibitory), based on the Zhuang-ABCA-1 transcriptomic dataset. However, real cortical connectivity differs dramatically between more refined cell types: pyramidal neurons and interneurons have distinct connection targets, and connectivity is strongly layer-specific. At the same time, we use a single global density parameter ρ to set the average number of connections per neuron across the entire cortex. A region-specific ρ could better capture the known variation in local synaptic density across cortical areas, for example the higher synaptic density of primary sensory regions relative to higher-order association areas. Future versions of the model could allow a region- and cell-type-specific ρ derived from region-level synapse-density atlases together with cell-type-resolved connectivity Harris et al. (2019), which would simultaneously address the limitations noted above.

Also, our model uses uniform synaptic weights within each synapse type: every AMPA synapse has conductance *g*_AMPA_ and every GABA synapse has conductance *g*_GABA_. In reality, synaptic strength varies with presynaptic and postsynaptic cell type, anatomical distance, and the distribution of synaptic weights is typically heavy-tailed (e.g. log-normal Buzsáki and Mizuseki (2014)). Extending our algorithm to sample synaptic weights from realistic distributions would be a natural next step and is likely necessary for quantitative comparisons with electrophysiological recordings.

### Cortical model and simulation

Compared to previous large-scale models Capone et al. (2023); Roberts et al. (2019), our model advances in incorporating the detailed spatial distribution of neurons in the 3-D computational domain. Without this information, spatiotemporal dynamics cannot be accurately analyzed and visualized. Recent research Pang et al. (2023); Gulbinaite et al. (2024) has highlighted the significant role that brain geometry plays in neuronal functions. In this work, we did not further use the spatial information of the cortex, such as the geometry of the cortical manifold, which would be an interesting direction for future work.

At the same time, although we are using one of the largest spatial transcriptomic datasets up to date, 300,000 neurons are still only a small portion of the total neuronal population in the mouse cortex. The number of neurons in a network will have important effects on aspects such as synaptic scaling. Among the total population, we only modeled two kinds of neurons, which is also a simplification. Therefore, enriching our model by including a higher number of different types of neurons and studying how they affect the conclusion in this paper is highly meaningful.

For simulations, we choose to randomly stimulate the total population of layer 4 neurons as a way to mimic subcortical input and generate traveling waves, which can be unrealistic. Subcortical structures, such as the thalamus, are vital to cortical dynamics like slow-wave activity Steriade et al. (1993) and are known to regulate traveling waves Ye et al. (2023). Therefore, a direct and important future improvement would be adding subcortical structures to the model. In this work, we choose two parameters that affect the dynamic regime of the network, which are the level of the stimuli controlled by the Poisson voltage bump and the coupling strength between neurons. Other parameters, including excitation-inhibition balance, are also likely to shape the dynamical regime Ahmadian and Miller (2021) and should be explored in future work.

Synaptic and conduction delays are essential to cortical dynamics such as traveling waves Petkoski and Jirsa (2022); Deco et al. (2009); Golomb and Ermentrout (2000). In our model, because the synapses are modeled by a first-order kinetic model (Neuronal and synaptic model), both excitatory and inhibitory synapses have a rising time that mimics the synaptic delay. However, we did not include conduction delay in our study. Conduction delay is thought to have a proportional relationship with the white matter path length of the connection between two neurons Lemaréchal et al. (2022). In our model, although we have the 3D positions of neurons, we do not have geo-metric information about the cortical manifold. Two neurons can be very close in the Cartesian coordinates measured by the Euclidean distance, but very far in terms of the length of the actual connection in the brain. Therefore, incorporating accurate conduction delay in the model is an important future direction.

### Quantitative measurements of 3-D traveling waves

There are many techniques available for identifying and measuring large-scale neuronal spatiotem-poral patterns Townsend and Gong (2018); Gutzen et al. (2024). While methods for detecting wave dynamics in three-dimensional data do exist Budzinski et al. (2023), most published algorithms are designed to analyze 2-D data, so our simulation data, which is intrinsically 3-D, presents new challenges for measurement. For instance, we need to identify neuron boundaries to exclude un-realistic boundary effects when calculating the phase gradient field (Quantitative measurement of neuronal activity). Additionally, working with high-resolution 3-D data is computationally expensive. Despite these challenges, we extended and verified the previous phase-based method to quantify the level of macroscopic wave activity in our simulations. However, as noted in the original work Rubino et al. (2006), phase gradient directionality only indicates planar wave activity and may fail to identify other wave types such as spiral waves, which are observed in the cortex Muller et al. (2016). We also did not analyze other cortical dynamics, such as sinks and sources, which have appeared in our simulation. Extending more sophisticated methods from 2-D to 3-D to identify diverse spatiotemporal patterns in experimental or simulated data will be a meaningful direction for future research.

### Comparing with experimental data

Across the paper, we have chosen the intracellular voltage of neurons as the primary data for the analysis. The voltage signals, though, do not directly resemble experimental measures such as local field potential (LFP), provide a quick way to analyze the overall dynamics of the whole network. There is strong experimental evidence of cortical spatiotemporal waves from measurements of LFP and multiunit activity reflecting extracellular electrical potentials in regions across the cortex Destexhe et al. (1999). Recent experimental studies of cortical wave patterns have achieved significant improvements in spatiotemporal resolution via methods such as wide-field calcium imaging Manley et al. (2024) and optical voltage imaging Liang et al. (2021). Our simulation framework allows many opportunities to compare with these experimental data.

However, one must be careful when comparing these simulations to experimental data, especially LFPs or other methods measuring extracellular potential rather than intracellular activity. Further work is needed to connect high-resolution mathematical models with experimental data based on the imaging techniques used. A more detailed quantitative comparison with experimental cortical-wave studies, such as the cortex-wide voltage-imaging data of Mohajerani et al. (2013) or the bidirectional visual-evoked waves reported by Aggarwal et al. (2022), is left as a future direction.

### Neural oscillation and traveling wave

Different neural oscillations are known to have different generating mechanisms. For example, the gamma oscillation is conjectured to be caused by coordinated interaction of excitatory and inhibitory neurons in the cortex Buzsáki and Wang (2012). And slow-wave activity is conjectured to be caused by thalamocortical interaction Steriade et al. (1993). In our work, we did not fully investigate how different kinds of oscillations emerge in the network from realistic mechanisms, but only studied how these oscillations produce different levels of macroscopic traveling waves in the cortex a posteriori. In the future, it is essential to combine the established understanding of mechanisms behind different oscillations with experiments to further verify the conclusions drawn in this work.

## Methods and Materials

### Use of the spatial transcriptomic data

The original spatial transcriptomic study Zhang et al. (2023) provides four datasets of cell information identified from the mouse brain. Among these, we selected the largest dataset, Zhuang-ABCA-1, which includes 2.8 million cells from one hemisphere, with approximately 2.6 million (N = 2,616,328) cells registered to the Allen Brain Atlas. Of these 2.6 million cells, about half are non-neuronal cells and were removed before constructing the connectivity. Since we are only modeling the glutamatergic and GABAergic neurons, we only kept these two kinds of neuronal populations, totaling approximately 1 million neurons (N = 1,050,440). As a result, our final dataset consists of N = 1,050,440 neurons (Fig. 1a) in one hemisphere. Within this population, the isocortex contains 297,812 neurons. The cortical connectivity is constructed by applying our algorithm to the total cortical population.

### Use of the voxelized connectivity data

The projection data we use in this study comes from Knox et al. (2018), which provides the connection strength between any pairwise voxel in the isocortex. The most important part of using the projection data is determining the number of connections between neurons in any pairwise voxels based on the connection strength. Let *W* represent the voxelized connectivity matrix, where *W*_*ab*_ represents the connection strength from voxel *a* to voxel *b*. In this work, we assume the number of connections between two voxels is proportional to the connection strength, meaning the number of connections from voxel *a* to voxel *b* is *N*_*ab*_ = λ*W*_*ab*_, where λ is a factor that will be calculated based on the density parameter.

Before building the connectivity, we first choose the density parameter, ρ, which is the average number of connections per neuron. The density parameter ρ is a free parameter that controls the trade-off between computational feasibility and biological fidelity: higher values of ρ yield more connections per neuron and thus higher biological realism, at the cost of greater memory and runtime Morrison et al. (2007). Let vector *x* represent the number of neurons in each voxel. If λ = 1, then the total number of connections in the whole connectivity is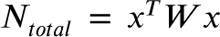, and the average number of connections per neuron is 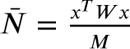, where *M* is the total number of neurons. Therefore, if we let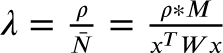, we will get the ideal average number of connections per neuron. Consequently, 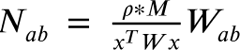. In the code, after calculating λ beforehand, we can multiply the whole projection matrix *W* by λ to derive a connection matrix, whose each entry now represents the number of connections between any two voxels.

### Construction and storage of the connectivity

Three types of connectivity are used in our simulations: Allen, local and uniform connectivity. Allen connectivity is constructed by applying our algorithm to the cortical neurons in Zhuang-ABCA1 based on the method described above. Uniform and local connectivity are constructed in a similar fashion: for each neuron, we first read out the number of outgoing connections in the corresponding Allen connectivity, then we randomly choose the same number of neurons from the total population without repetition for the uniform connectivity, or the same number of nearest neighbors for local connectivity. In this way, the out-degree distribution of all three connectivity will be the same, making them more comparable. In particular, local connectivity is constructed through nearest-neighbor coupling, where we use the K-nearest neighbors (KNN) algorithm Cover and Hart (1967) to choose a fixed number of outgoing connections to nearest neighbors for each neuron based on spatial position. For all three connectivity, self-connection is not allowed.

When running our connectivity algorithm to build the Allen connectivity, a large number of neurons can pose some challenges to our connectivity algorithm. To tackle this challenge, we loop through the voxels for a significant speed-up. As a result, our algorithm will take *O*(*M N*) time, where M is the number of neurons that contain all the neurons and N is the number of voxels. N has a maximum since the computational domain is finite, and in the end, this algorithm is about two orders of magnitude faster than iterating through every neuron.

In our model, we choose ρ = 300, which means there are about 90 million synapses in the model since we have 297,812 neurons. To store such huge connectivity, we use a compressed row form representation of the connectivity matrix, which is also used in similar packages Dai et al. (2020).

### Neuronal and synaptic model

We use a Hodgkin-Huxley type neuron model that can produce different firing behaviors of a cortical neuron Pospischil et al. (2008). Each neuron has four ionic channels: voltage-gated sodium, voltage-gated potassium, slow non-inactivating potassium, and leak channels. The pyramidal neurons will have all four ionic channels, while the inhibitory neurons do not have the slow non-inactivating potassium channels because they are fast-firing and do not usually exhibit firing adaptation. Detailed equations of each current and the change of gating variables are provided in the appendix. The parameters of the neuronal model are also drawn from the original paper and presented in Table S1 in the appendix.

For synapses, we model both glutamatergic (AMPA) or GABAergic synapses in our cortical network. Both synapses are modeled by a conductance-based, first-order kinetic model. Please see the supplement for more details.

### Simulation

To simulate the computational model, we have to solve two sets of equations at every time step: the Hodgkin-Huxley equations for the electrophysiology model and the conductance-coupled synaptic equations. We solve the Hodgkin-Huxley equations using an established implicit leapfrog method DeWoskin et al. (2015), allowing for large timesteps due to increased stability. Both electrophysiology and synaptic equations are solved in parallel on GPU in our framework. The overall parallelism is based on the neuron, which means each set of Hodgkin-Huxley equations of a neuron is put up on an individual thread of GPU to solve at every timestep. Similarly, at any time step, we first check for all neurons firing and then assign each thread to process each of the firing neurons’ downstream connections. Please see the supplements for a detailed description of the numerical methods.

Each simulation lasts for 3 seconds, and we only record the last 1 second of the simulation to avoid the transient at the beginning of the simulation. The initial conditions of all neurons are randomly chosen at the beginning of each simulation. To be more specific, the initial voltage of all neurons is drawn from a uniform distribution between −65 mV and −5 mV, and all gate variables are drawn from a uniform distribution between 0.02 and 0.05. For each simulation, we collect the voltage traces of all neurons with a sampling frequency of 2000 Hz.

In all simulations, the total excitatory population of layer 4 neurons is stimulated with a Poisson spike train at a rate of 10 Hz, delivering a fixed-amplitude voltage bump *V*_stim_ on each Poisson event. To adjust the level of drive we sweep *V*_stim_ from 5 to 32.5 mV in steps of 2.5 mV. To adjust the coupling strength we sweep the excitatory conductance *g*_AMPA_ from 0.005 to 0.050 nS in steps of 0.005 nS, with the inhibitory conductance always fixed at 10% of the excitatory conductance. The full *g*_AMPA_ ×*V*_stim_ grid (10 × 12 = 120 simulations per connectivity) is used for the Poisson-stimulation analyses in Figs. 4-6.

### Quantitative measurement of neuronal activity

To quantify spatiotemporal dynamics, we extend a phase-based method Rubino et al. (2006) that has primarily been used for 2-D data but naturally generalizes to 3-D (figure Supplement 1 illustrates the full pipeline on a representative snapshot). Because the Hilbert transform is most interpretable when applied to narrowband signals, the simulated voltage traces are first interpolated onto a regular Cartesian grid that follows the Allen Brain Institute common coordinate framework (CCFv3), then band-pass-filtered into a frequency band, and only then converted to a generalized phase via the Hilbert transform. Concretely:

1. Spatial interpolation. We construct a regular 3-D Cartesian grid following the discretization of the Allen Brain Institute common coordinate framework (CCFv3), of size 132 × 80 × 114 at 100 μm spatial resolution, and linearly interpolate the per-neuron voltage time series onto this grid; grid cells that contain no neurons are masked out and excluded from later steps.
2. Band-pass filtering. The gridded voltage time series are bandpass-filtered into the five canonical bands (delta 0.5-4 Hz, theta 4-8 Hz, alpha 8-12 Hz, beta 12-30 Hz, gamma 30-100 Hz) using a fourth-order Butterworth filter applied with zero-phase filtfilt.
3. Generalized phase. For each narrowband-filtered gridded signal, we apply the Hilbert transform and extract the band-specific generalized phase Φ(*x*, *t*) for that band.
4. Phase gradient directionality. We compute the phase gradient ∇Φ on the 3-D grid using a finite-difference scheme and use it to compute the phase gradient directionality (PGD) *P* (*t*) Rubino et al. (2006), our quantitative measure of macroscopic wave activity:

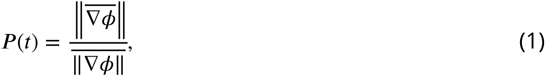

where the horizontal bar denotes the spatial average across the grid. Higher PGD means the gradient field is more aligned across spatial locations, indicating a more coherent planar wavefront.

In computing the gradient field, the boundary grid cells that contain no neurons are excluded by masking, implemented with the PyVista package Sullivan and Kaszynski (2019); both interpolation and gradient computation use the intrinsic routines of the 3-D Visualization Toolkit (VTK) Schroeder et al. (1998).

The Kuramoto order parameter *R*(*t*) = *e*^*i*Φ(*x,t*)^ _*x*_ is computed by averaging *e*^*i*Φ^ across the valid (neuron-containing) grid cells of the same per-band phase field, and reduced to its maximum over each 1-s recording window for the synchrony curves in **Figure 5**.

The same per-band pipeline is run independently for each of the five bands in the per-band analyses (Figure 4c, **Figure 5**, **Figure 6**, figure Supplement 2-figure Supplement 5, figure Supplement 6, figure Supplement 1, figure Supplement 2). The LFP analyses in **Figure 6** use the same pipeline applied to the per-neuron synaptic-LFP estimates (see LFP estimation from intracellular voltage) after interpolation onto the same grid.

### LFP estimation from intracellular voltage

To connect our simulated intracellular voltage traces to experimentally measured local field potentials (LFP), we used a post-hoc pipeline that reconstructs the synaptic current received by every neuron directly from the voltage traces and the connectivity used in the simulation Telenczuk et al. (2020). Since LFP primarily reflects the net transmembrane synaptic current in a local population Destexhe et al. (1999), using the reconstructed synaptic current as an LFP proxy is a natural and widely adopted choice in large-scale simulations.

Concretely, for each neuron *i* at time *t* we first compute the sigmoidal neurotransmitter output used in the simulation’s synaptic model,

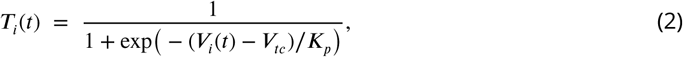

with *V*_*tc*_ = 2 mV and *K*_*p*_ = 5 mV matching the parameters used in the simulation. We then use the CSR connectivity matrix to propagate this presynaptic output to the postsynaptic neurons, keeping excitatory and inhibitory sources separate. Let *A*^*E*^ and *A*^*I*^ denote the incoming-connection adjacency matrices restricted to excitatory and inhibitory presynaptic neurons, respectively (obtained by transposing the outgoing CSR matrix and masking columns by the presynaptic E/I label); the excitatory and inhibitory synaptic activation at neuron *j* is

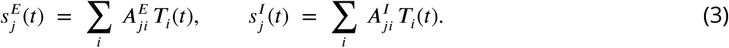

The net synaptic current per neuron is then

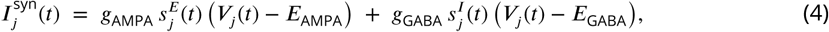

with reversal potentials *E*_AMPA_ = 0 mV, *E*_GABA_ = −70 mV, and the same conductances *g*_AMPA_, *g*_GABA_ used in the corresponding simulation. This per-neuron current *I*^syn^(*t*) plays the role of an LFP-like proxy: spatially averaging *I*^syn^ over a local population yields a signal analogous to an extracellularly measured LFP. The pipeline is implemented with sparse matrix multiplications and streams the voltage traces in chunks, so it scales to the full 297,812-neuron population across the 1-second recording window.

We then run the same 3-D phase-gradient pipeline described in Quantitative measurement of neuronal activity on the reconstructed LFP. The per-neuron synaptic current is interpolated onto the Allen Brain Atlas 3-D grid, bandpass-filtered into the delta, theta, alpha, beta, and gamma bands, converted to a phase field via the Hilbert transform, and used to compute PGD.

### Visualization

Throughout the paper, we visualize neurons by coloring them based on their intracellular voltage. The visualization is straightforward for a single neuron such as in figure Supplement 1, but in **Figure 2**, **Figure 4** and **Figure 6**, we also need to calculate the average voltage of 300 neurons nearby. This is done by identifying the closest 300 neighbors for each neuron using the K-Nearest-Neighbor algorithm and calculating the average. All visualizations are made using the PyVista package Sullivan and Kaszynski (2019).

## Videos

**Video 1.** Macroscopic wave activity emerging in the Allen-connectivity Poisson simulation of **Figure 2**. Two paired three-dimensional renderings of the same simulation are animated side-by-side over a 500 ms window of the recorded activity (1 ms cadence, 30 fps playback). Left: 300-nearest-neighbour local-mean intracellular voltage (the smoothed view of Figure 2b). Right: raw single-neuron intracellular voltage with no spatial averaging (the single-cell-resolution view of figure Supplement 1), showing how the macroscopic wavefront emerges from the underlying spiking activity. Both panels use the turbo colormap. The trace at the bottom is the global mean voltage of all ∼300,000 cortical neurons over the full 1 s recording, with the red dashed band marking the 50 ms snapshot window highlighted in Figure 2b and a vertical cursor tracking the current animation frame. Simulation parameters match **Figure 2**: Allen connectivity, *g*_AMPA_ = 0.045 nS, *V*_stim_ = 12.5 mV, 10 Hz Poisson spike train delivered to all layer-4 excitatory neurons.

**Video 2.** Theta-band traveling waves under Allen, local and uniform connectivity, corresponding to the three simulations of **Figure 4**. The three brain panels show the theta-band (4-8 Hz) generalized phase at single-neuron resolution for three matched simulations that share the same Poisson stimulation protocol but differ only in connectivity: Allen (left), local (center) and uniform (right).

**Video 3.** The macroscopic wavefront is preserved in the LFP estimate, supplementing Figure 6a. Two paired renderings of the same Allen Poisson simulation are animated side-by-side across a 200 ms window of the recorded activity (1 ms cadence, 30 fps playback). Left: 300-nearest-neighbour local-mean intracellular voltage on the turbo colormap with *V* ∈ [−65, −50] mV. Right: 300-nearest-neighbour local-mean post-hoc-reconstructed synaptic-LFP estimate on the diverging RdBu colormap, *z*-scored across the rendered window.

## AI Usage

Generative AI was used during revision to assist with language editing; all scientific content, analyses, interpretations, and final wording were reviewed and approved by the authors.

## Data availability

Raw data related to this study, including the neuronal and connectivity data, along with the code and simulation data related to the figures, have been uploaded to the following repository: **Error! Hyperlink reference not valid.**

## Acknowledgments

We thank Ningyuan Wang for a helpful discussion about work and coding. This work is dedicated to Steven Brown, who provided many helpful suggestions.

We acknowledge the following funding: ARO MURI (Multidisciplinary University Research Initiatives) W911NF-22-1-0223, HFSP (Human Frontier Science Program) RGP0019/2018, and NSF DMS (National Science Foundation Division of Mathematical Sciences) 2052499. The views and conclusions contained in this document are those of the authors and should not be interpreted as representing the official policies, either expressed or implied, of the Army Research Office or the U.S. Government. The U.S. Government is authorized to reproduce and distribute reprints for Government purposes, notwithstanding any copyright notation herein. Any opinions, findings, and conclusions or recommendations expressed in this material are those of the author(s) and do not necessarily reflect the views of the National Science Foundation.

---

**Figure 1—figure supplement 1.**
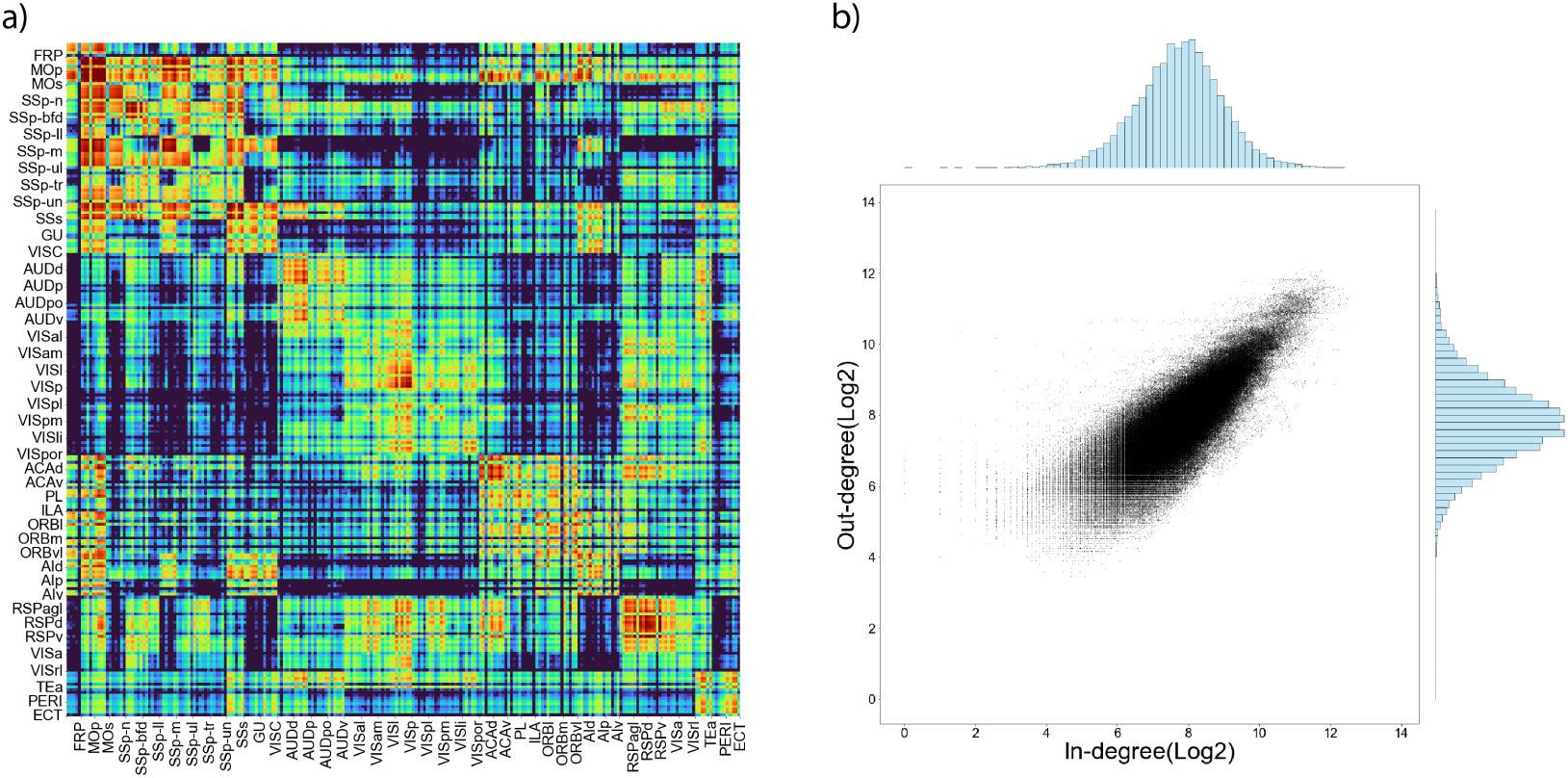
Connectivity matrix with detail region legend and degree distribution. a The same connectivity matrix shown in Fig. 1c, but with detailed region legend. b The out and in-degree distribution of all neurons in log 2 scale.

**Figure 2—figure supplement 1.**
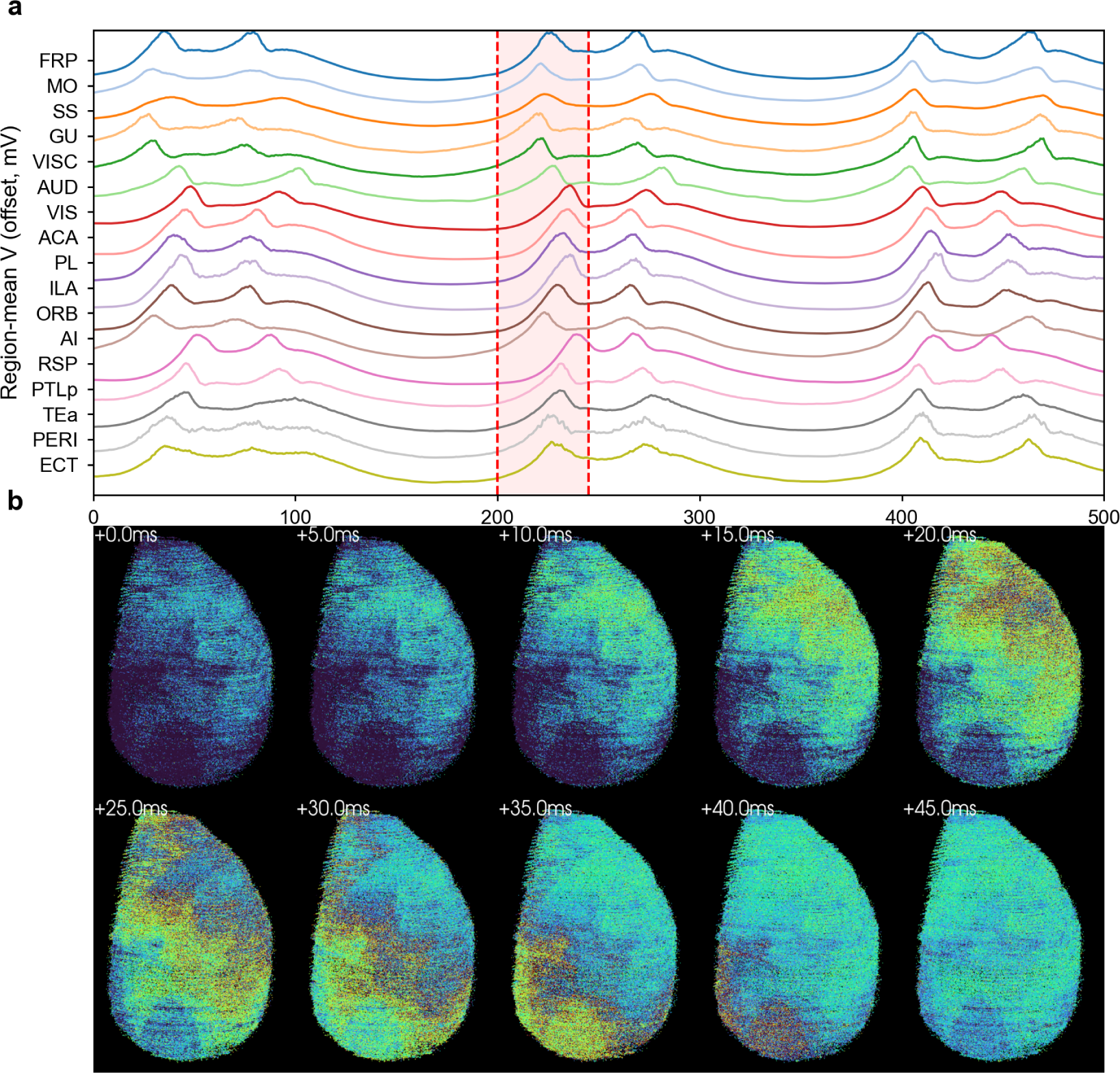
Single-neuron-resolution view of the simulation in Fig. 2. (a) Region-mean voltage traces (offset for clarity), one trace per major cortical region in anterior-to-posterior order. (b) Two rows of five voltage snapshots taken every 5 ms over the window highlighted in (a), showing the wavefront propagation at single-neuron rather than KNN-averaged resolution.

**Figure 3—figure supplement 1.**
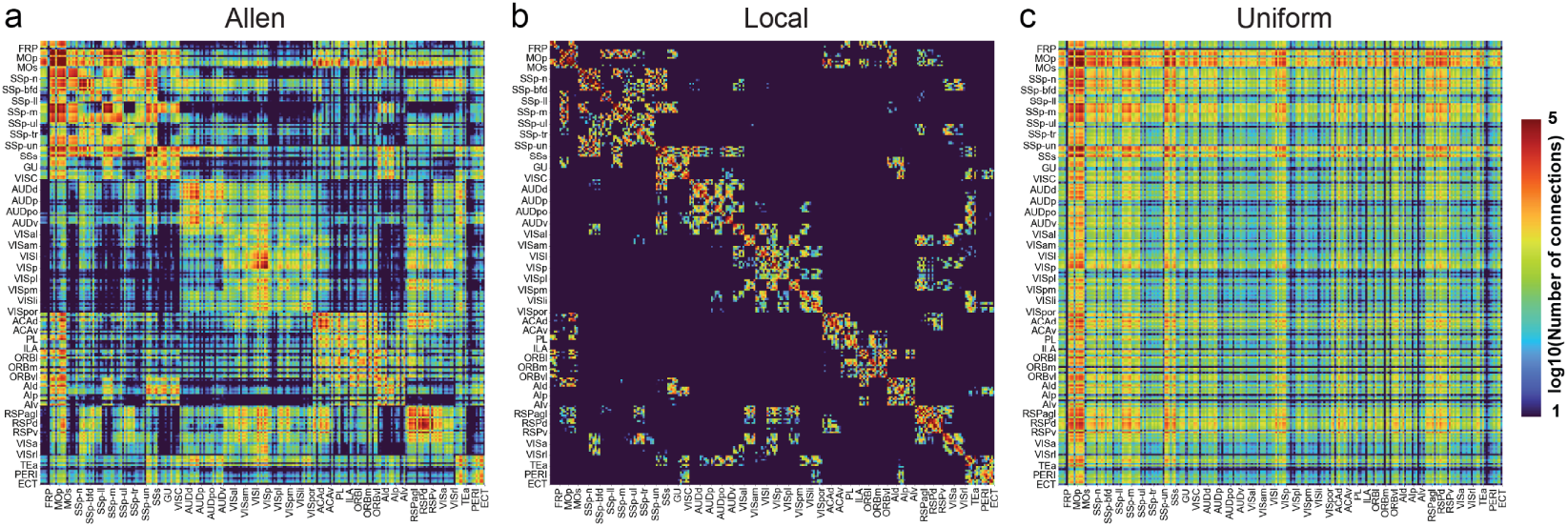
Comparison of the three connectivity matrices: a Allen, b local and c uniform connectivity. For local connectivity, the diagonal terms represent the connections within the same region. And the off-diagonal terms exist because these regions are physically adjacent to each other.

**Figure 4—figure supplement 1.**
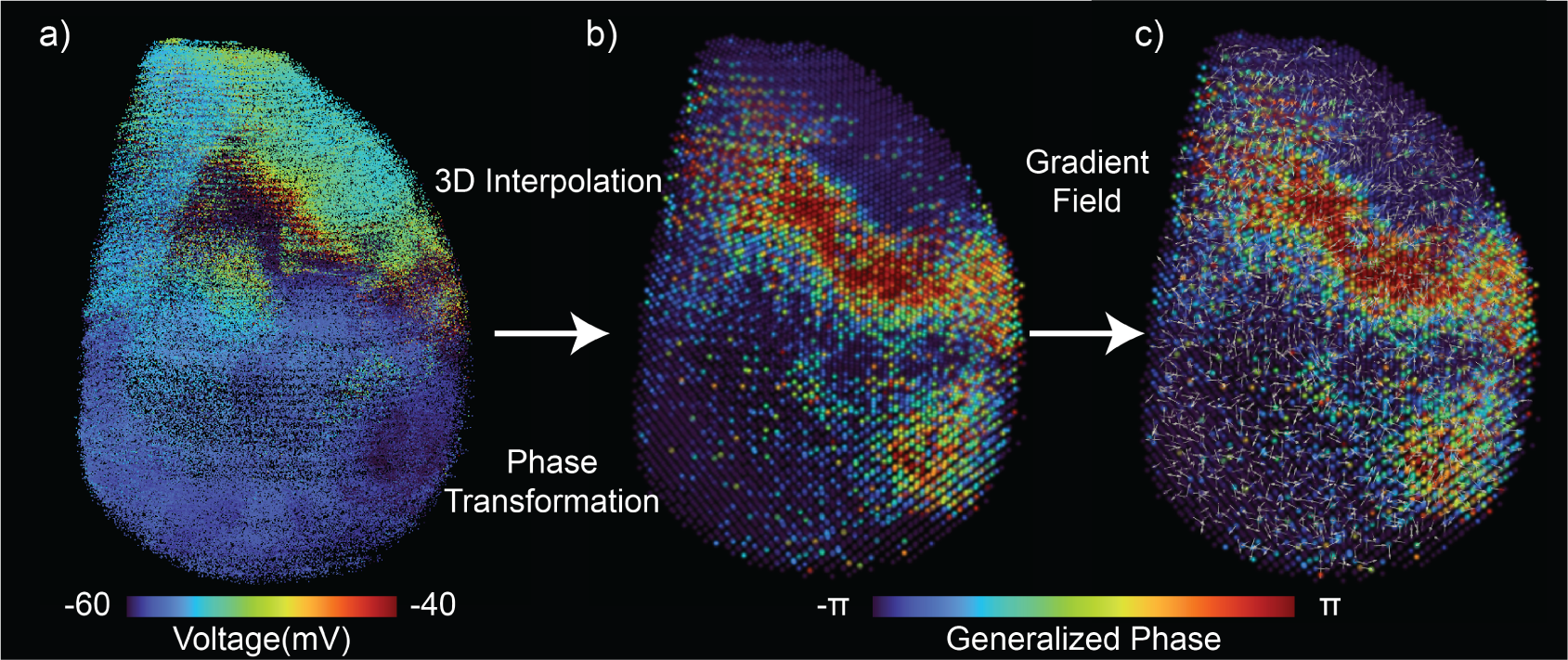
Illustration of the three-dimensional phase-gradient pipeline used to quantify macroscopic wave activity (Quantitative measurement of neuronal activity), shown for a single representative snapshot. (a) The per-neuron intracellular voltage is linearly interpolated onto the regular CCFv3 Cartesian grid (color: membrane voltage in mV). (b) After band-pass filtering and the Hilbert transform, every grid point carries a generalized phase Φ(*x*, *t*) (color: phase from − π to π). (c) The spatial gradient ∇Φ of the phase field is computed at each grid point (white arrows); the phase gradient directionality (PGD) summarizes how well aligned these gradient vectors are and is our scalar measure of macroscopic traveling-wave activity.

**Figure 4—figure supplement 2.**
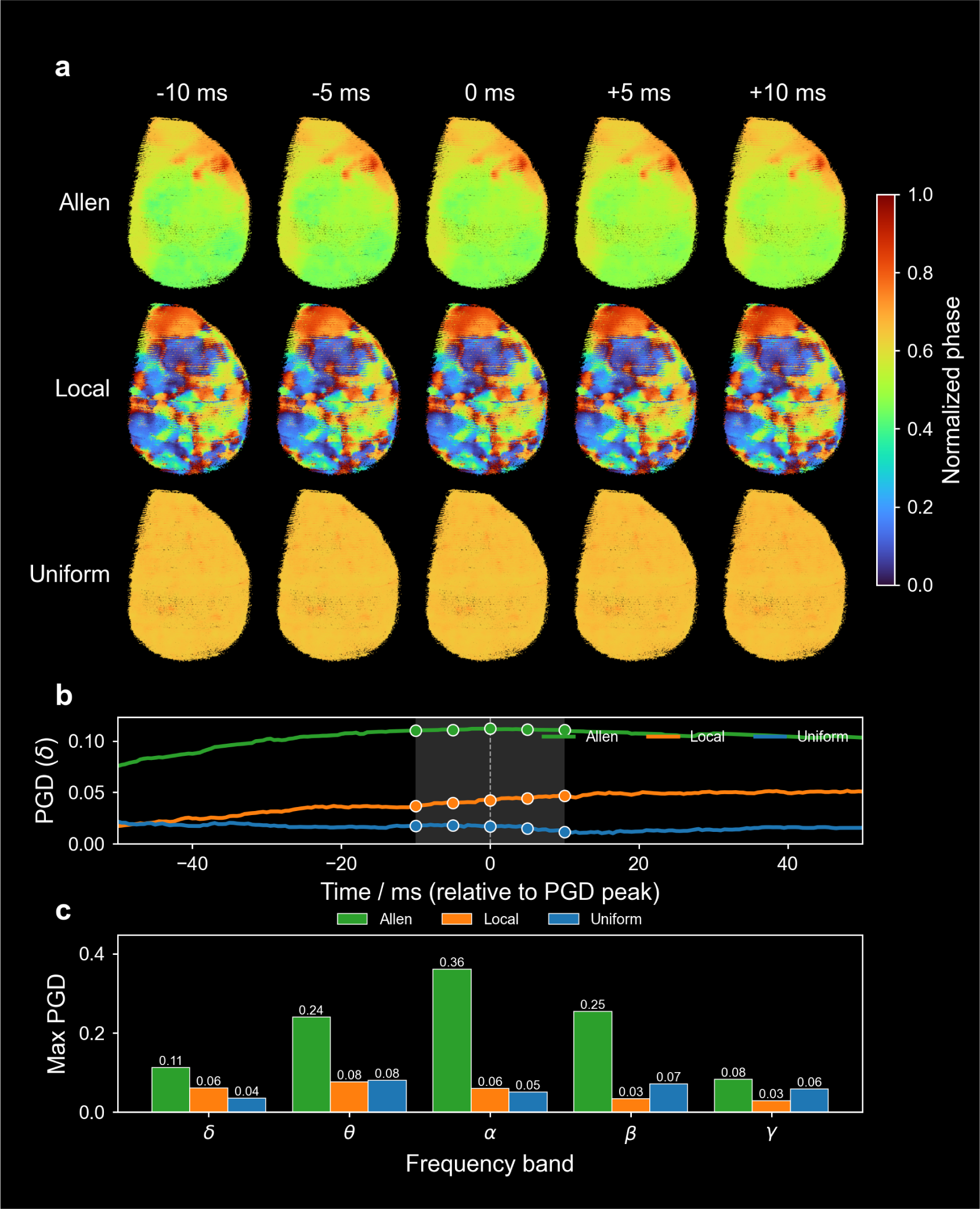
delta-band (0.5-4 Hz) version of Figure 4 (which shows the theta band). (a) 3 × 5 spatial snapshots of the delta-band generalised phase (rows: Allen / Local / Uniform; columns: five frames centered on the Allen-PGD peak); phase is computed on the voltage signal with 300-NN spatial smoothing in the complex (Hilbert) domain, with no Cartesian-grid interpolation. (b) delta-band PGD(*t*) over a 100 ms window centered on the Allen peak, all three connectivities. (c) Per-band max PGD bars for the same simulation (identical across the four band figures, included to match the main-text Figure 4 layout).

**Figure 4—figure supplement 3.**
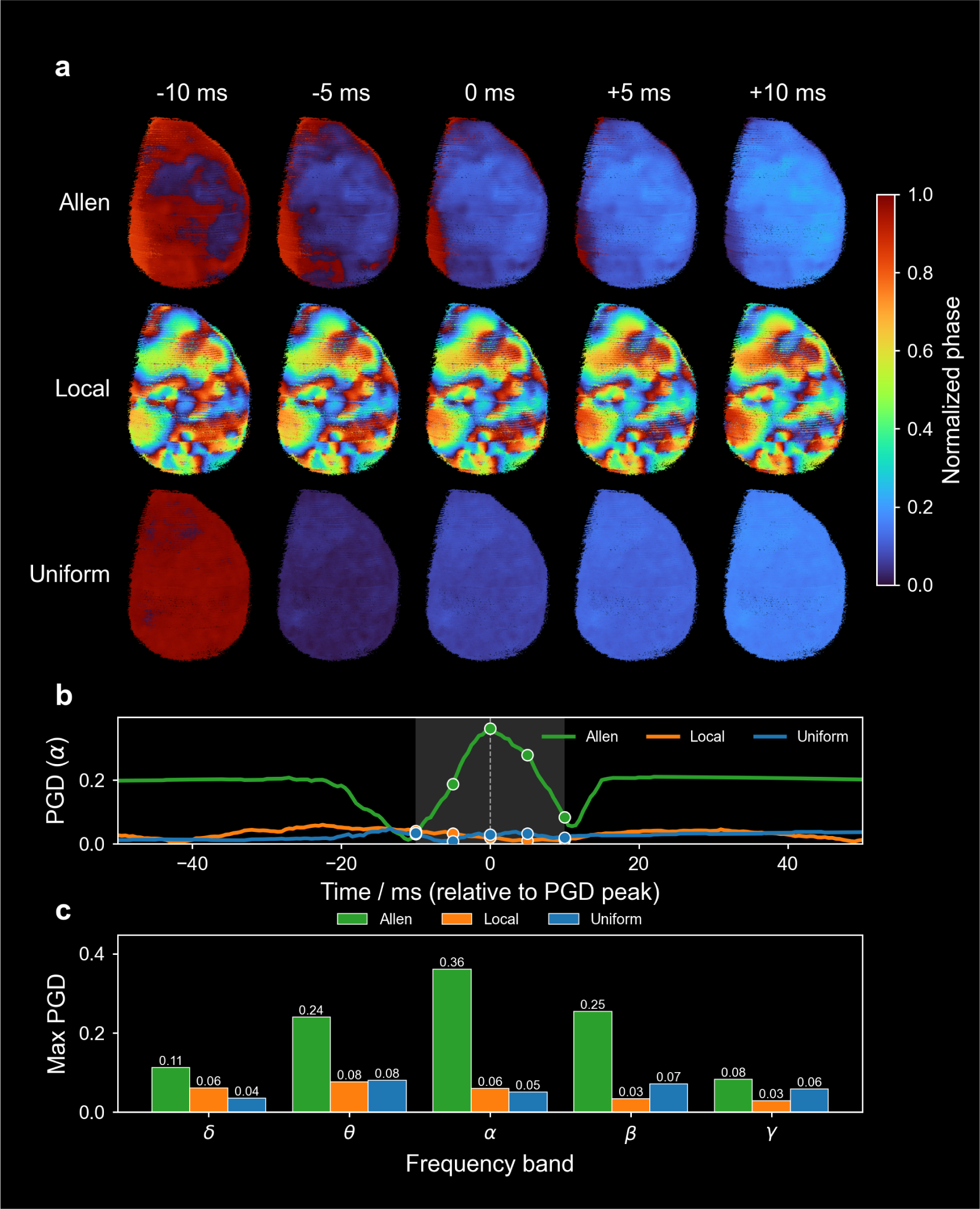
alpha-band (8-12 Hz) version of Figure 4; layout and conventions as in figure Supplement 2.

**Figure 4—figure supplement 4.**
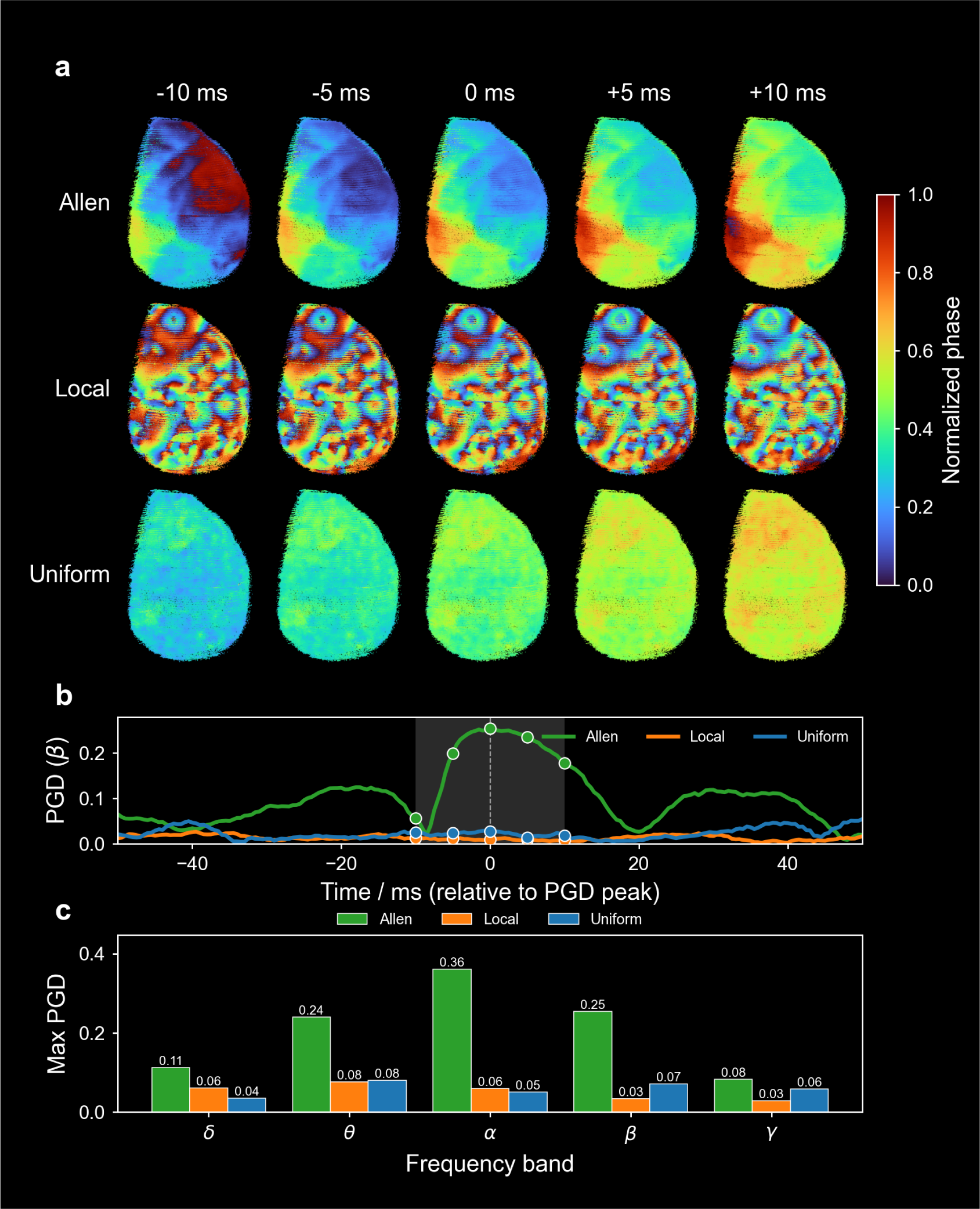
beta-band (12-30 Hz) version of Figure 4; layout and conventions as in figure Supplement 2.

**Figure 4—figure supplement 5.**
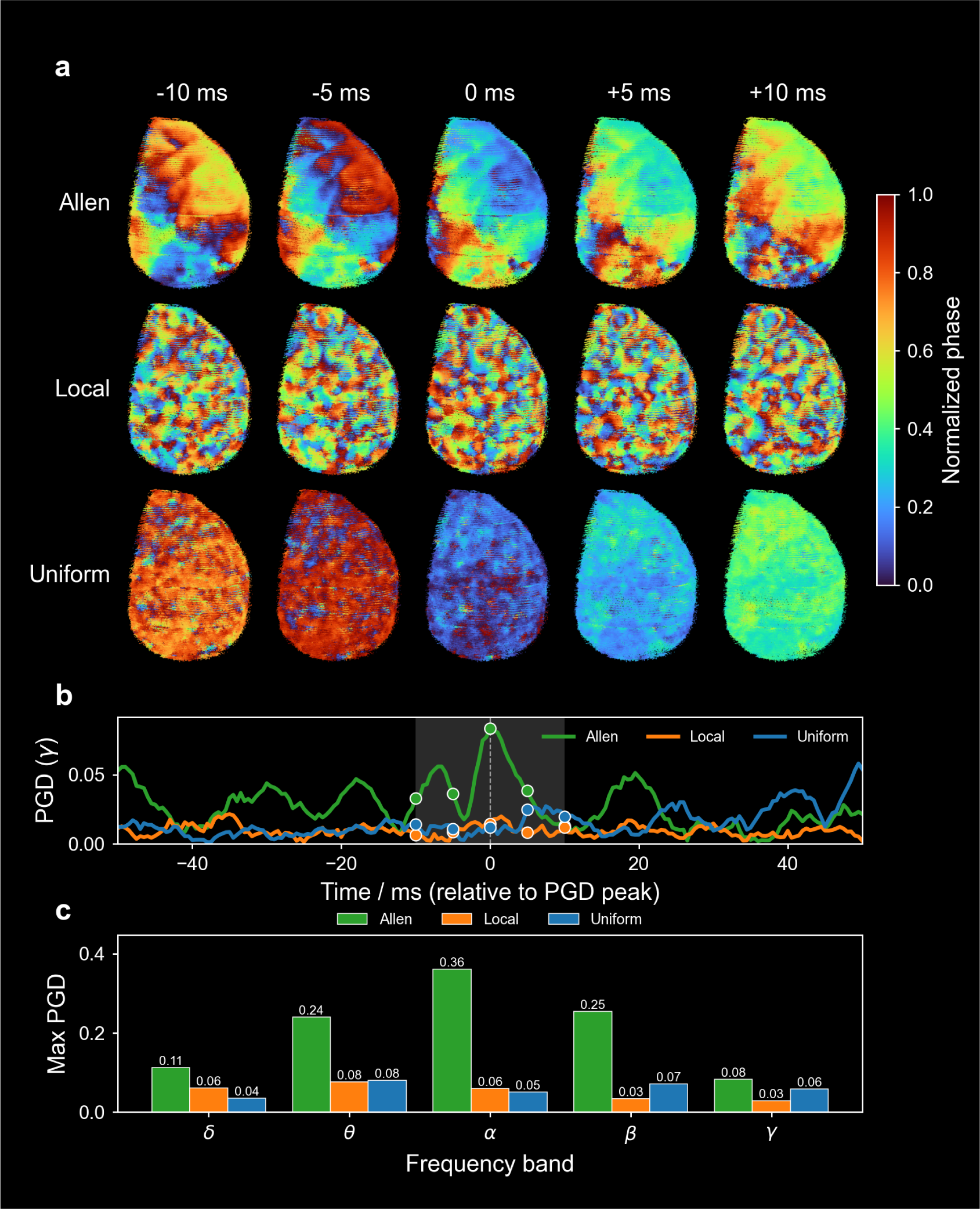
gamma-band (30-100 Hz) version of Figure 4; layout and conventions as in figure Supplement 2.

**Figure 4—figure supplement 6.**
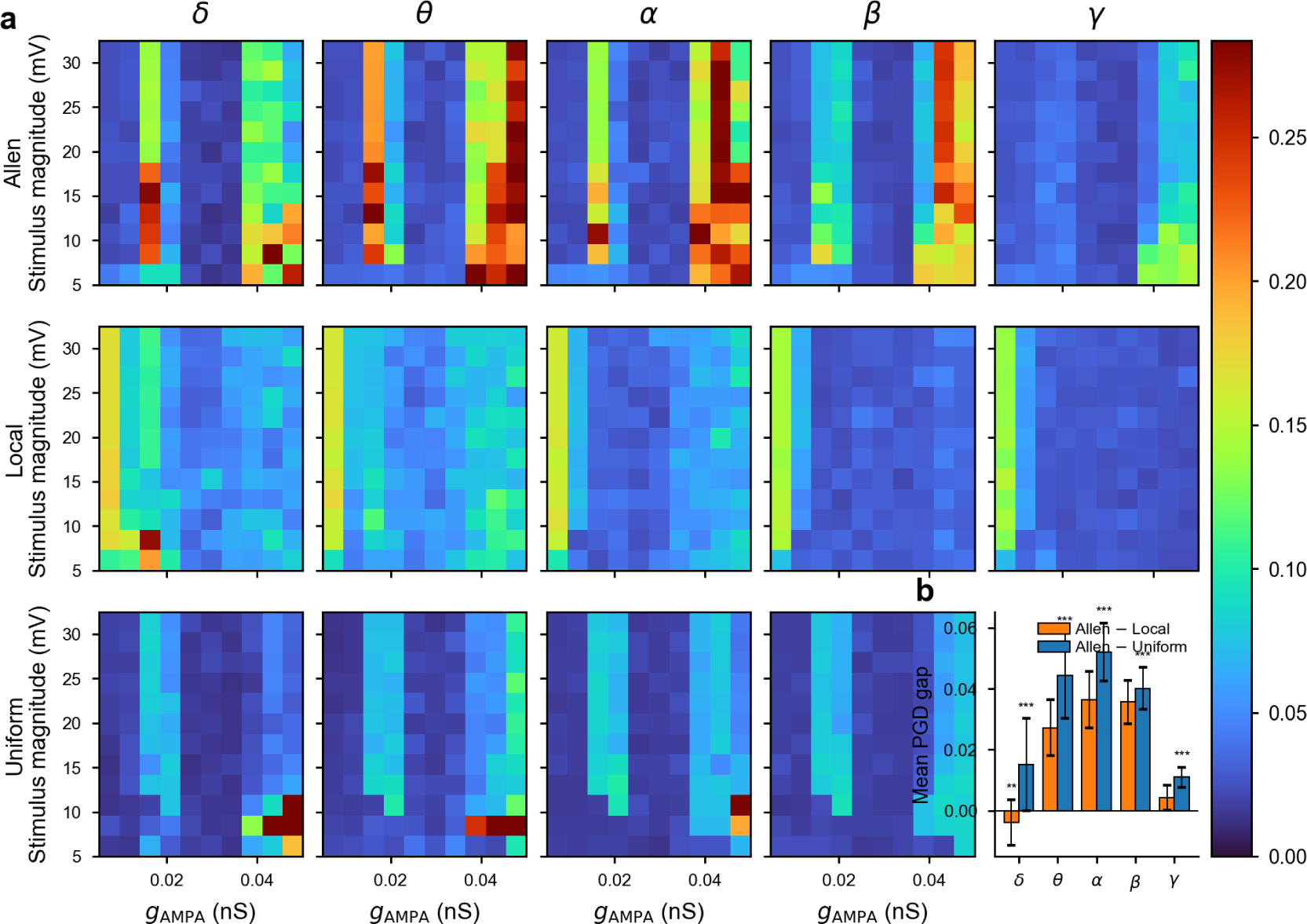
Per-band PGD heatmaps and Allen advantage across the Poisson (*g*_AMPA_, *V*_stim_) sweep. (a) 3 × 5 grid of max-PGD heatmaps over the Poisson (*g*_AMPA_, *V*_stim_) sweep; rows are the three connectivities (Allen / Local / Uniform), columns the five frequency bands (delta, theta, alpha, beta, gamma). Every heatmap shares the same axes (*g*_AMPA_ on *x*, *V*_stim_ on *y*) and the turbo colour scale on the right (Max PGD). (b) Per-band Allen-minus-opponent mean PGD gap (mean ± SEM across the 10 × 12 grid), with grouped bars for Allen − Local (orange) and Allen − Uniform (blue) and two-sided significance stars (*: *p* < 0.05; **: *p* < 0.01; ***: *p* < 10^−4^); panel (b) occupies the bottom-right slot of the 3 × 5 grid.

**Figure 5—figure supplement 1.**
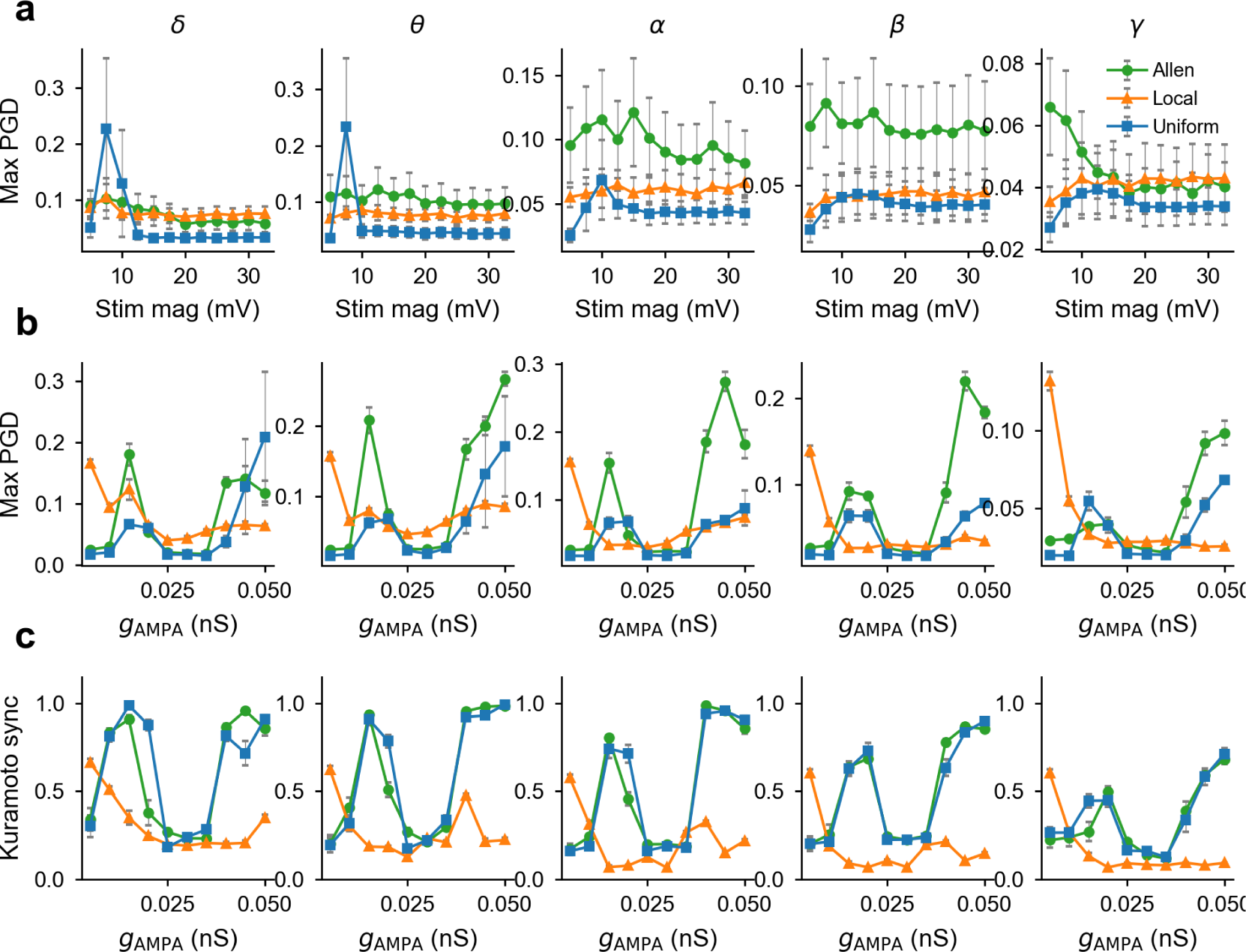
Per-band versions of Figure 5c-e. A 3 × 5 grid combining (a) max PGD versus *V*_stim_ (averaged over the *g*_AMPA_ sweep; supplement to Figure 5c); (b) max PGD ver-sus *g*_AMPA_ (averaged over the *V*_stim_ sweep; supplement to Figure 5d); and (c) maximum Kuramoto synchrony *R*(*t*) = *e*^*i*Φ(*x,t*)^ _*x*_ versus *g*_AMPA_ (averaged over the *V*_stim_ sweep; supplement to Figure 5e). Columns are the five frequency bands (delta, theta, alpha, beta, gamma); error bars are SEM across the orthogonal sweep parameter. The peak-dip-recovery coupling profile of Figure 5d and the matching synchrony dip are visible in every band for Allen and uniform connectivity; local connectivity decays monotonically across all bands.

**Figure 5—figure supplement 2.**
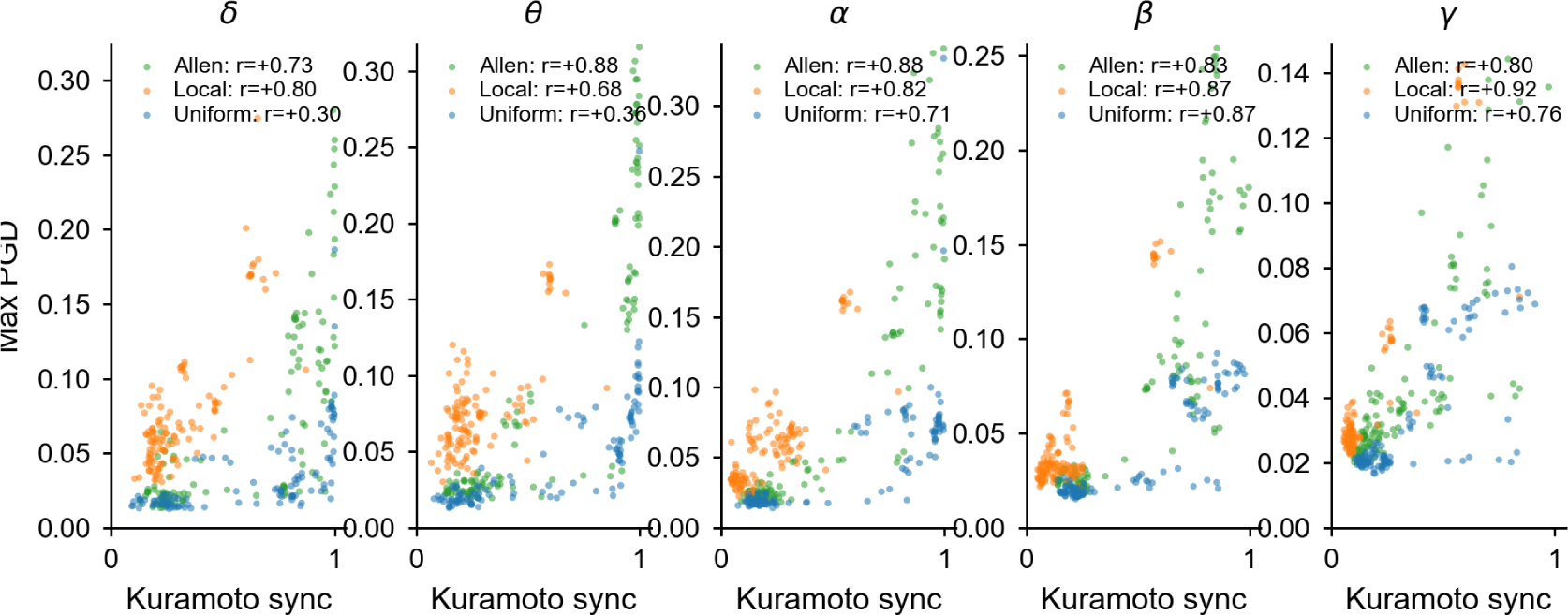
Synchrony-PGD scatter across all bands. One point per (*g*_AMPA_, *V*_stim_) combination on the 10 × 12 sweep grid, with *x* = max Kuramoto synchrony and *y* = max PGD, coloured by connectivity (Allen / Local / Uniform). Per-connectivity Pearson *r* values are shown in each panel’s legend. Synchrony and PGD are positively correlated in every band and every connectivity (all *p* < 10^−3^). This extends Figure 5f from the alpha band to all five canonical bands.

